# Multimodal single-cell/nucleus RNA-sequencing data analysis uncovers molecular networks between disease-associated microglia and astrocytes with implications for drug repurposing in Alzheimer’s disease

**DOI:** 10.1101/2020.09.23.310466

**Authors:** Jielin Xu, Pengyue Zhang, Yin Huang, Lynn Bekris, Justin Lathia, Chien-Wei Chiang, Lang Li, Andrew A. Pieper, James B. Leverenz, Jeffrey Cummings, Feixiong Cheng

## Abstract

Systematic identification of molecular networks in disease relevant immune cells of the nervous system is critical for elucidating the underlying pathophysiology of Alzheimer’s disease (AD). Two key immune cell types, disease-associated microglia (DAM) and disease-associated astrocytes (DAA), are biologically involved in AD pathobiology. Therefore, uncovering molecular determinants of DAM and DAA will enhance our understanding of AD biology, potentially identifying novel therapeutic targets for AD treatment. Here, we present an integrative, network-based methodology to uncover conserved molecular networks between DAM and DAA. Specifically, we leverage single-cell and single-nucleus RNA sequencing data from both AD transgenic mouse models and AD patient brains, drug-target networks, metabolite-enzyme associations, and the human protein-protein interactome, along with large-scale patient data validation from the MarketScan Medicare Supplemental Database. We find that common and unique molecular network regulators between DAM (i.e, *PAK1, MAPK14*, and *SYK*) and DAA (i.e., *NFKB1, FOS*, and *JUN*) are significantly enriched by multiple neuro-inflammatory pathways and well-known genetic variants (i.e., *BIN1*) from genome-wide association studies. Further network analysis reveal shared immune pathways between DAM and DAA, including Fc gamma R-mediated phagocytosis, Th17 cell differentiation, and chemokine signaling. Furthermore, integrative metabolite-enzyme network analyses imply that fatty acids (i.e., elaidic acid) and amino acids (i.e., glutamate, serine, and phenylalanine) may trigger molecular alterations between DAM and DAA. Finally, we prioritize repurposed drug candidates for potential treatment of AD by agents that specifically reverse dysregulated gene expression of DAM or DAA, including an antithrombotic anticoagulant triflusal, a beta2-adrenergic receptor agonist salbutamol, and the steroid medications (fluticasone and mometasone). Individuals taking fluticasone (an approved anti-inflammatory and inhaled corticosteroid) displayed a significantly decreased incidence of AD (hazard ratio (HR) = 0.858, 95% confidence interval [CI] 0.829-0.888, *P* < 0.0001) in retrospective case-control validation. Furthermore, propensity score matching cohort studies also confirmed an association of mometasone with reduced incidence of AD in comparison to fluticasone (HR =0.921, 95% CI 0.862-0.984, *P* < 0.0001).

## Introduction

Alzheimer’s disease (AD) is a devastating neurodegenerative disease and it is estimated that it will affect 16 million Americans and 90 million people worldwide by 2050^1^. The incidence of AD is expected to double by 2050^2^. The attrition rate for AD clinical trials (2002-2012) is estimated at 99.6%^3^ and improved methods of drug discovery and development are needed. There are multiple risk factors implicated in disease pathogenesis, such as genetic factors, local and systemic inflammation, psychosocial stress responses, and many other unknown factors^4^. The underlying genetic basis and molecular mechanisms of disease pathobiology/physiology remain under investigation. Furthermore, predisposition to AD involves a complex, polygenic, and pleiotropic genetic architecture^5^ . The traditional reductionist paradigm (‘one gene, one drug, one disease’) overlooks the inherent complexity of human diseases and has often led to treatments that are inadequate or accompanied by adverse effects^6^. Given the heterogeneous clinical presentation, AD is no longer considered a neuronal-centric disease; recent studies strongly implicate a crucial role of neuro-inflammation in the pathobiology of AD^7^. Broad anti-inflammatory therapies have not been clinically efficacious against AD, suggesting a pressing need to better understand the heterogeneity of these immune cells and identify drug targets for novel treatment development.

Advances in single-cell technologies are beginning to uncover crucial roles of the immune systems in disease onset and the pathogenesis of AD. Recent single-cell/nucleus RNA-sequencing (scRNA-seq or snRNA-seq) studies have suggested essential roles for microglia and astrocytes, such as determining the “normal” and pathological immune cell subpopulations in AD^8, 9^. For example, disease-associated microglia (DAM) was identified as a unique microglia subtype associated with AD pathogenesis^8^. Disease associated astrocytes (DAA) have been identified in early stage of AD and become more abundant with AD progression^10^. Cytokines, the primary immune messenger, can mediate astrocytes to influence the microglial activation state (e.g., *CCL2* and *ORM2*) and help microglia modulate astrocytic phenotypes and functions (e.g., *IL-1α* and *TNF-α*)^11^. A growing body of evidence suggests that both microglia and astrocytes are exquisitely sensitive to their environment that can be affected by the dysregulation of multiple biochemical pathways, such as abnormal lipid metabolism, in AD pathogenesis^12^. Systematic identification of the underlying molecular mechanisms between DAM and DAA would advance understanding of disease biology and offer potential drug targets for novel therapeutic development in AD.

Existing data resources, including genomics, transcriptomics, and interactomics (protein-protein interactions [PPIs]), have not yet been fully exploited to understand the causal disease pathways in AD^13^. Integrative analyses of genomics, transcriptomics, and other omics enable us to elucidate the cascade of molecular events contributing to complex neuro-inflammatory mechanisms, including microglia and astrocytes. This will accelerate the translation of high-throughput omics findings to innovative therapeutic approaches for AD by integrating knowledges from both microglia and astrocytes. In this study, we propose an integrative multi-omics, network-based methodology to identify novel underlying molecular determinants for DAM and DAA in AD. Specifically, we systematically characterized the molecular networks for both microglia and astrocytes by incorporating large-scale snRNA-seq and scRNA-seq data into the human protein-protein interactome. We showed that the identified DAM or DAA specific molecular networks offer novel pathobiological pathways and potential drug targets for AD. We demonstrated that drugs reversing the dysregulated gene expression of DAM or DAA offer potential treatment strategies for AD and we validated these agents in a large-scale, real-world patient database.

## Results

### Network-based methodology pipeline

In this study, we presented an integrative multi-omics, network-based methodology to uncover molecular networks of DAM and DAA and to prioritize drug candidates for potential treatment of AD by reversing dysregulated gene expression in DAM and DAA. We integrated scRNA-seq and snRNA-seq data from both AD transgenic mouse models and AD patients brain tissues, drug-target networks, enzyme-metabolite associations, the human protein-protein interactomes, along with large-scale patient database validation (**Figure 1**). The whole procedure is divided into 4 components: i) We first collected the 4 recent sc/snRNA-seq datasets (**Supplementary Table 1**) covering both microglia and astrocytes from either AD transgenic mouse models or human AD brains; ii) We performed standard sc/snRNA-seq data analysis (**Methods**) which includes quality control, cell/nucleus clustering and differentially expressed genes (DEGs) analysis in sequential order for each sc/snRNA-seq profile; iii) We built the molecular network for DAM and DAA using the state-of-the-art network-based algorithm by integrating sc/snRNA-seq data into the human protein-protein interactome (**Methods**); iv) We prioritized repurposed drugs for potential treatment of AD by identifying those that specifically reverse dysregulated gene expression of microglia and astrocytes: if drug-induced up- or down-related genes are significantly enriched in the dysregulated molecular networks of DAM or DAA, these drugs will be prioritized as potential candidates for treatment of AD. Finally, top drug candidates were validated further using the state-of-the-art pharmacoepidemiologic observations of a large-scale, real-world patient database (**Figure 1**).

**Figure 1.**
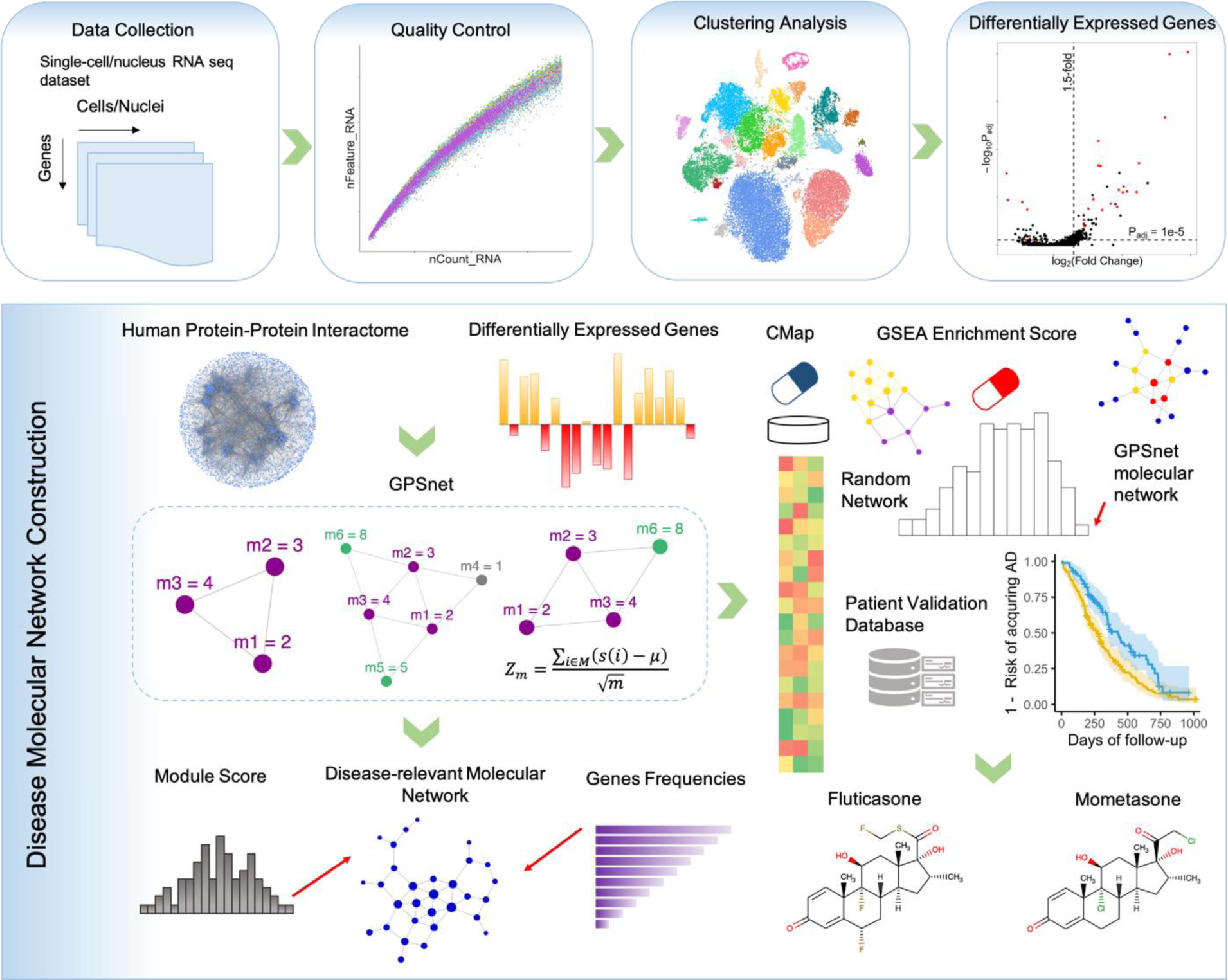
A diagram illustrating the network-based framework. A standard single cell / nucleus RNA-sequencing data analysis pipeline includes quality control, clustering analysis and differentially expressed genes (DEGs) analysis. We built the molecular network using the state-of-the-art network-based algorithm by integrating sc/snRNA-seq data into the human protein-protein interactome (**Methods**). Next, we prioritized repurposed drugs for potential treatment of AD by identifying those that specifically reverse dysregulated gene expression for microglia and astrocyte: if drug-induced up- or down-related genes are significantly enriched in the dysregulated molecular networks, these drugs will be prioritized as potential candidates for treatment of AD. Finally, top drug candidates were validated further using a large-scale, real-world patient database.

### Discovery of disease-associated microglia specific molecular networks

We compared expression of cell marker genes (*CST7, LPL, P2RY12*, and *CX3CR1*) for DAM among all cell/nucleus clusters for snRNA-seq (**Figure 2A, B**) and scRNA-seq (**Supplementary Figures 1**) profiles, respectively. Here, we used homeostasis-associated microglia (HAM^14^) as control groups. We discover that, under the snRNA-seq profile, the DAM cells have much higher abundance (88%, normalized nucleus abundance percentage) in 5XFAD mice compared to wild-type (WT) mice (12%, **Table 1A** and **Supplementary Figure 2A**). Yet, the normalized nucleus abundance percentages of HAM cells (33%) in 5XFAD mice is lower than WT mice (67%, **Table 1A** and **Supplementary Figure 2A**). Similarity, when considering the scRNA-seq profile, the normalized cell abundance percentage of the DAM in 5XFAD mice (94%) is much higher than WT mice (6%, **Table 1B** and **Supplementary Figure 2B**) as well. And the corresponding normalized cell abundance percentages of HAM cells in 5XFAD mice (47%) is marginally lower than WT mice (53%, **Table 1B** and **Supplementary Figure 2B**) counted by the scRNA-seq profile. Altogether, both sc/snRNA-seq profiles show significantly elevated abundance of the DMA in 5XFAD mice compared to WT mice.

**Figure 2.**
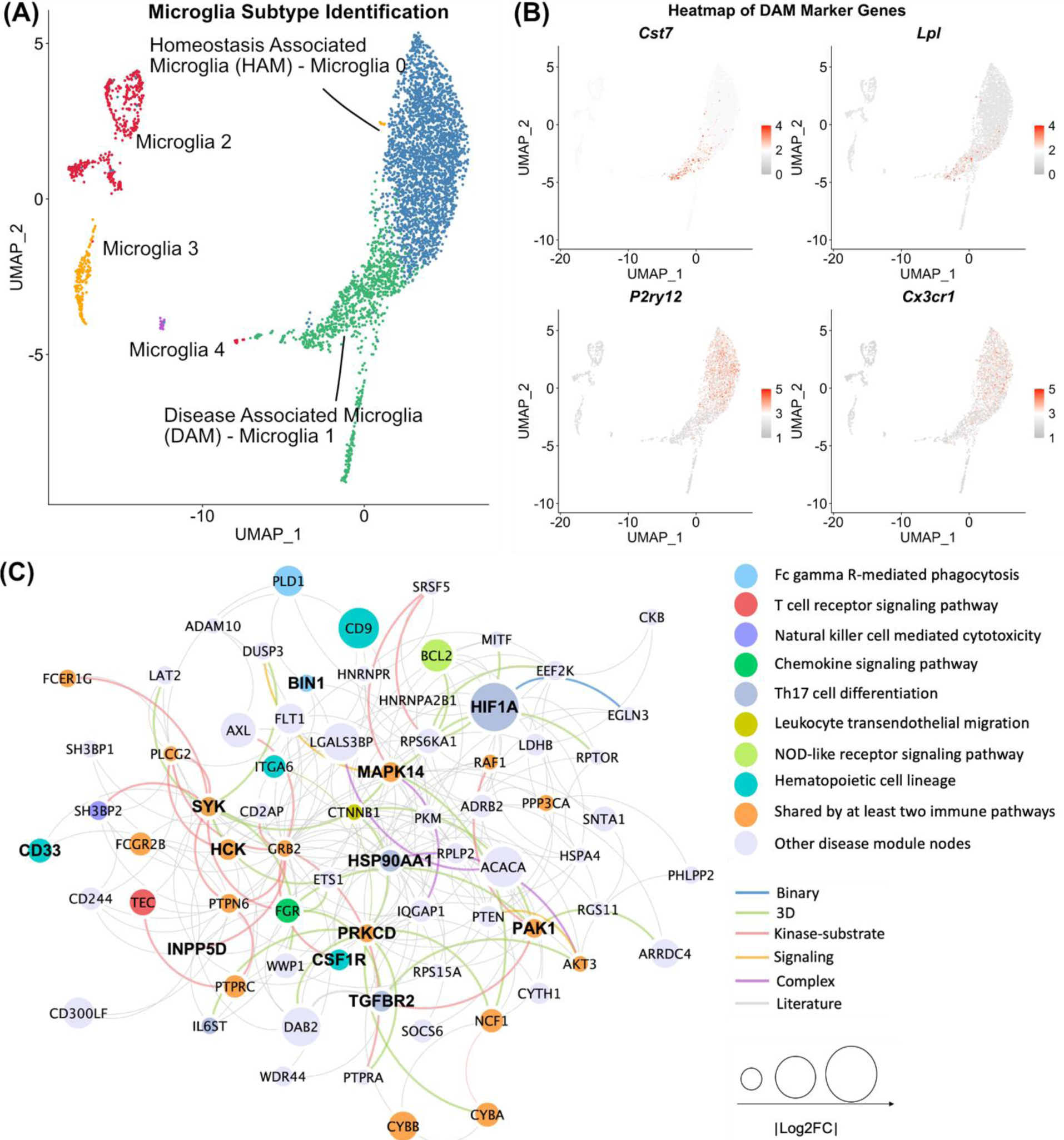
Discovery of disease-associated microglia (DAM) specific molecular networks for the AD transgenic mouse model. (**A**) Uniform manifold approximation and projection (UMAP) plot of clustering 4,389 microglia cells: blue cluster denotes the homeostasis associated microglia (HAM) and green cluster denotes the DAM. (**B**) Expression levels (heatmap) of representative marker genes (up-regulation in DAM: *Cst7* and *Lpl* and down-regulation in DAM: *P2ry12* and *Cx3cr1*) in different microglia sub-clusters. (**C**) A predicted DAM specific molecular network contains 227 protein-protein interactions (PPIs) connecting 72 genes/proteins. Node sizes are proportional to their corresponding |log2FC| during differential expression analysis. Nodes are color coded by known KEGG immune pathways. Edges are color coded by different experimental evidences of PPIs (**Method**).

We further examined genome-wide differential expression analyses of DAM compared to HAM in 5XFAD mice as shown in Volcano plots (**Supplementary Figure 3**). As expected, several key AD genes or microglia markers are significantly differentially expressed in DAM compared to HAM in 5XFAD mice, including *APOE, TREM2, CST7, PYRY12*, and *CX3CR1* (**Supplementary Figure 3**). To identify novel molecular pathways underlying DAM, we systematically searched molecular networks using our recently published network-based approach, called GPSnet^15^. Specifically, GPSnet re-constructs the network module using a selected seed gene (i.e., differentially expressed genes [DEGs]) from the PPIs, and each time expands the module by adding a qualified candidate neighboring gene that could improve the module score measured by the fold changes of differential expression analysis. The final molecular network is constructed by aggregating modules with top ranked genes that frequently appear in top ranked modules (**Methods**). The identified molecular networks for DAM using snRNA-seq (termed snDAMnet) and scRNA-seq (termed scDAMnet) datasets are shown in **Figure 2C** and **Supplementary Figure 1C**. The snDAMnet includes 227 PPIs connecting 72 unique genes (e.g., *BIN1, HCK, HSP90AA1, IL6ST, PAK1, PRKCD*, and *SYK*, **Supplementary Table 1**). Myc box-dependent-interacting protein 1 (*BIN1),* a well-established risk gene for AD by the International Genomics of Alzheimer’s Project, contains a microglia-specific enhancer and promoter by a genome-wide significant AD variant rs6733839^16^. We next collected AD-associated genes from multiple sources, including expert curated repositories, GWAS catalog^17^, animal models and the scientific literatures as described in a previous study^18^. We found that genes in snDAMnet were significantly enriched in 31 AD-associated genes (adjusted p-value [q] = 1.75 x10^-10^, Fisher’s exact test, **Supplementary Table 1**), such as *ADAM10, BIN1, CD33*, and *MAPK14* (**Supplementary Table 1**). The scDAMnet contains 69 genes (e.g., *AXL, CST7, LYN, MERTK*, and *PYRY12,* **Supplementary Table 2**) connecting 97 PPIs. As expected, scDAMnet covers 27 AD-associated genes^18^ (e.g., *APOE, CCL3, CTSD, INPP5D,* and *MARCKS*, q = 5.00x10^-8^, Fisher’s exact test, **Supplementary Table 2**). We further found that genes in DAMnets are significantly enriched in multiple immune pathways (**Supplementary Tables 1** and **2**) as well. We observed that most of them were critical immune modulators related to AD (**Figure 2C**, **Supplementary Figure 1C,** and **Supplementary Table 1**). We next discussed the selected genes in snDAMnet and scDAMnet across 4 selected immune pathways: Fc gamma R-mediated phagocytosis, chemokine signaling pathway, Th17 cell differentiation, and hematopoietic cell lineag*e* (**Supplementary Tables 1** and **2**).

#### Fc gamma R-mediated phagocytosis

In total, we identified 15 genes (such as *BIN1, PRKCD, SYK, INPP5D*, and *HCK*) in the Fc gamma R-mediated phagocytosis pathway which were enriched in either snDAMnet or scDAMnet (**Supplementary Tables 1** and **2**). *BIN1* is one of the most important loci for late onset Alzheimer’s disease (LOAD)^16^. Several studies uncovered crucial functions of *PRKCD* in AD: a) A*β* stimulated protein kinase C delta type (*PRKCD*) to phosphorylate myristoylated alanine-rich C-kinase substrate (*MARCKS*) in microglia^19^ and phosphorylation of *MARCKS* was observed in microglia within plaques^20^; and b) inhibition of *PRKCD* reverse A *β* levels^21^. Spleen tyrosine kinase (*SYK*) has been shown to play a role in AD pathological lesions, and *SYK* was therefore considered as a potential drug target for AD^22^. Phosphatidylinositol 3,4,5-trisphosphate 5-phosphatase 1 (*INPP5D*), identified as one of the genetic risk factors for LOAD^23^, affects AD pathology by regulating microglia^24^. Inhibiting tyrosine-protein kinase (*HCK*) has proved to disturb microglia function and exacerbate neuropathology and neuroinflammation^25^.

#### Chemokine signaling pathway

Chemokine signaling is enriched in both snDAMnet and scDAMnet and are related with 13 genes, including *PAK1, CCL3, CCL4*, *CCR5 and LYN* (**Supplementary Tables 1** and **2**). Serine/threonine-protein kinase (*PAK1*) is dysregulated in AD and targeting the PAK signaling pathway offers a therapeutic strategy for treating AD^26^. C-C motif chemokine 3 and 4 (*CCL3* and *CCL4*) and C-C chemokine receptor type 5 (*CCR5*)^27^ have been shown to be upregulated in adult human microglia or in mouse microglia that were stimulated with A*β*. A recent study observed elevated activity of tyrosine-protein kinase (*LYN*) in AD patients, and inhibiting *LYN* expression prevents A*β*-induced neuronal cell death, suggesting *LYN* as an potential therapeutic target for AD^28^.

#### Th17 cell differentiation

We identified 6 genes (including *MAPK14, HIF1A, TGFBR2*, and *HSP90*) in the *Th17 cell differentiation* pathway (**Supplementary Table 1**). Reduced A*β* pathology was observed in mitogen-activated protein kinase 14 (*MAPK14*) -/- APP-PS1 transgenic AD mouse neurons, suggesting that inhibiting *MAPK14* could serve as a potential alternative to mitigate pathologies in neurons for AD^29^. The transcriptional factor hypoxia-inducible factor 1-alpha (*HIF1A*) is recognized as a key gene for a variety of neurodegenerative diseases, including AD, Parkinson’s disease (PD), and Huntington’s disease (HD)^30^. Insufficient levels of TGFBRs (TGF-beta receptors) are major risk factors of AD, and increasing TGF-beta receptor type-2 (*TGFBR2*) levels is a potential therapeutic strategy for AD^31^. HSP90 (heat shock protein 90), a chaperone protein, regulated tau pathology by forming macromolecular complexes with co-chaperones and inhibiting HSP90 mitigated tau pathology by proteasomal degradation^32^.

#### Hematopoietic cell lineage

Four genes (i.e., *CSF1R*, *CD33, CD9* and *ITGA6*) identified from the snDAMnet are involved in regulating the hematopoietic cell lineage pathway (**Supplementary Table 1**). Inhibiting macrophage colony-stimulating factor 1 receptor (*CSF1R*) in APP/PS1 mice reverses microglia from an inflammatory to an anti-inflammatory phenotype, suggesting that inhibiting *CSF1R* could treat microglia activation and AD^33^. Myeloid cell surface antigen *CD33*, is elevated in AD brain and could compromise the ability of microglia to remove A*β* plaques, suggesting it could serve as a possible therapeutic target for AD^34^. We did not find strong AD-related evidence for another two genes (*CD9* and *ITGA6*), revealing novel candidate genes that required further functional validation.

In summary, we identified that DAM specific molecular networks were significantly enriched in multiple AD-related immune pathways. Importantly, a variety of proteins in DAM specific molecular networks are profoundly involved in AD pathogenesis and offer potential drug targets for AD.

### Discovery of disease-associated astrocyte specific molecular networks

We compared gene expression of 11 DAA cell markers (*GFAP, CD44, HSPB1, SLC1A2*, and *PTN*) among all nuclei clusters and located non-disease associated astrocyte (non-DAA) and DAA clusters for the mouse snRNA-seq profiles (**Figures 3A** and **3B**). We found that a normalized nucleus abundance percentage of DAA cells in 5XFAD mice (99%) is significantly higher than WT mice (1%, **Table 1C** and **Supplementary Figure 2C**) in human snRNA-seq profiles. For the non-DAA cells, a normalized nucleus abundance percentage in 5XFAD mice (41%) is slightly lower than WT mice (59%, **Table 1C** and **Supplementary Figure 2C**). The human snRNA-seq profiles contains AD brain samples from 2 regions – entorhinal cortex (EC) and superior frontal gyrus (SFG). T-distributed stochastic neighbor embedding^35^ (TSNE) plots of DAA and non-DAA nuclei are presented in **Figure 4A** and **4B** for each brain region, respectively. Gene expression of 11 DAA cell markers (*GFAP, CD44, HSPB1, SLC1A2*, and *PTN*) among all nuclei clusters in both human brain regions are presented in **Supplementary Figure 4**. Volcano plots of DEGs (i.e., *GFAP, APOE* and *MYOC*) are shown for both DAA and non-DAA nuclei in either 5XFAD mice (**Supplementary Figure 5**) or human AD samples (**Supplementary Figure 6**). The molecular networks for mouse snRNA-seq (mDAAnet) and human snRNA-seq across two specific brain regions (hDAAECnet and hDAASFGnet), are shown in **Figures 3C** and **Figure 4C** and **4D**, separately. The mDAAnet includes 371 PPIs connecting 98 unique genes (e.g., *CD44, CTSD, ICAM1, MARCKS, NFKB1,* and *VCAM1*, **Supplementary Table 3**). It contains 39 AD-associated genes collected from DisGeNET^18^ (e.g., *CDH2, CLU, CTSD, FOS,* and *TGFBR2*, q = 1.70x10^-12^, Fisher’s exact test, **Supplementary Table 3**). The hDAAECnet contains 43 PPIs connecting 26 genes (e.g., *APC, DCLK2, ID3, PRKCA,* and *TNC*, **Supplementary Table 4**), including 11 AD-associated genes (e.g., *ATXN1, FGF2, HSP90AA1,* and *JUN*, q = 1.96x10^-3^, Fisher’s exact test, **Supplementary Table 4**). The hDAASFGnet contains 22 PPIs connecting 13 genes (e.g., *DCLK2, FOS,* and *TNC*, **Supplementary Table 4**), including 8 AD-associated genes (e.g., *FGFR3, FOS, HSP90AA1,* and *JUN*, q = 9.50x10^-4^, Fisher’s exact test, **Supplementary Table 4**).

**Figure 3.**
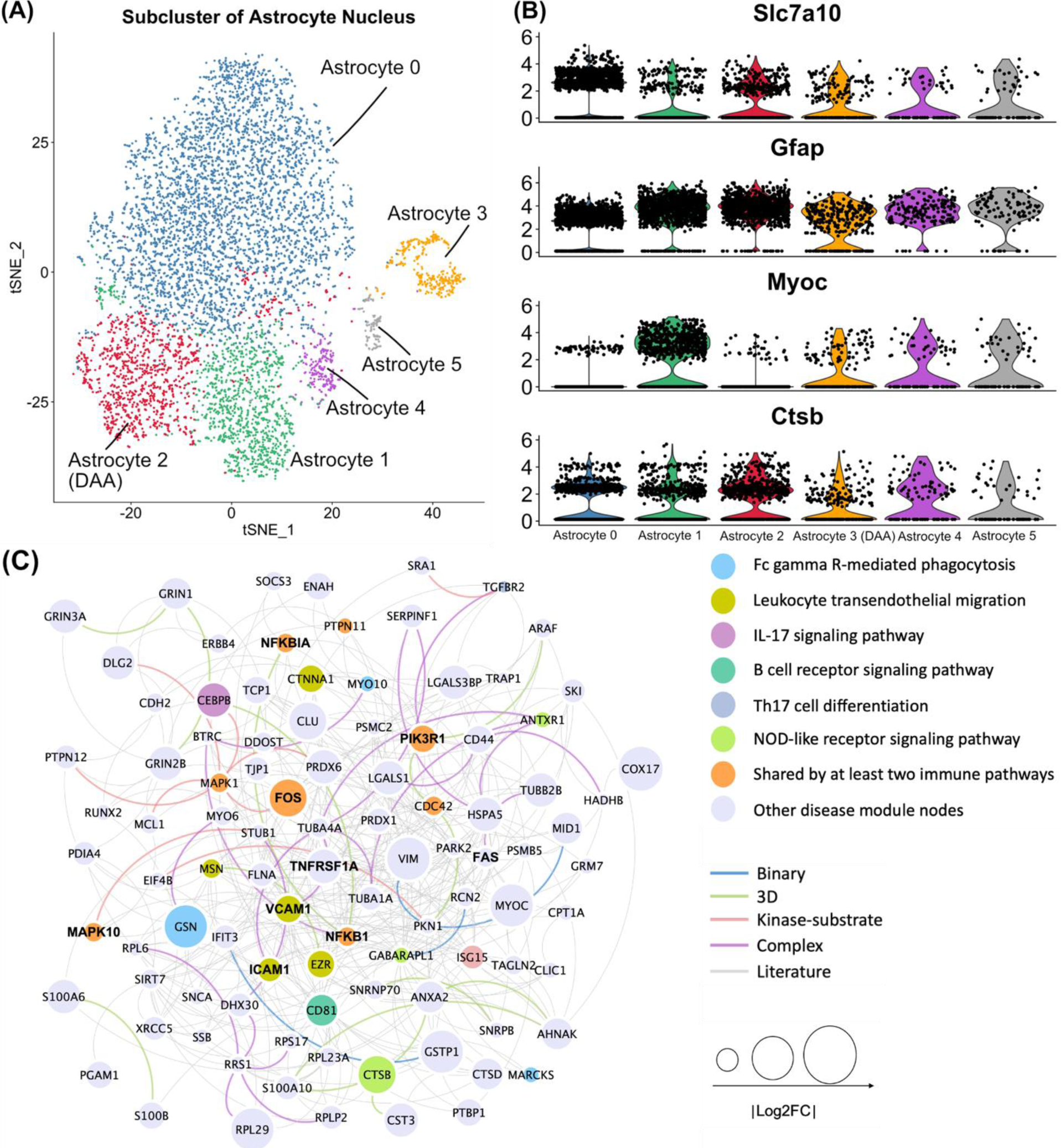
Discovery of disease-associated astrocyte (DAA) specific molecular networks in AD transgenic mouse model. (**A**) T-distributed stochastic neighbor embedding (TSNE) plot of clustering 7546 astrocyte nuclei. Red cluster denotes the disease associated astrocyte (DAA); (**B**) Stacked violin plot displaying the expression patterns of 9 representative genes across different astrocyte sub-clusters; (**C**) A predicted DAA specific molecular network contains 371 protein-protein interactions (PPIs) connecting 98 genes/proteins. Node sizes are proportional to their corresponding |log2FC| during differential expression analysis. Nodes are color coded by KEGG immune pathways, and edges are color coded by different experimental evidences of PPIs (see **Methods**).

**Figure 4.**
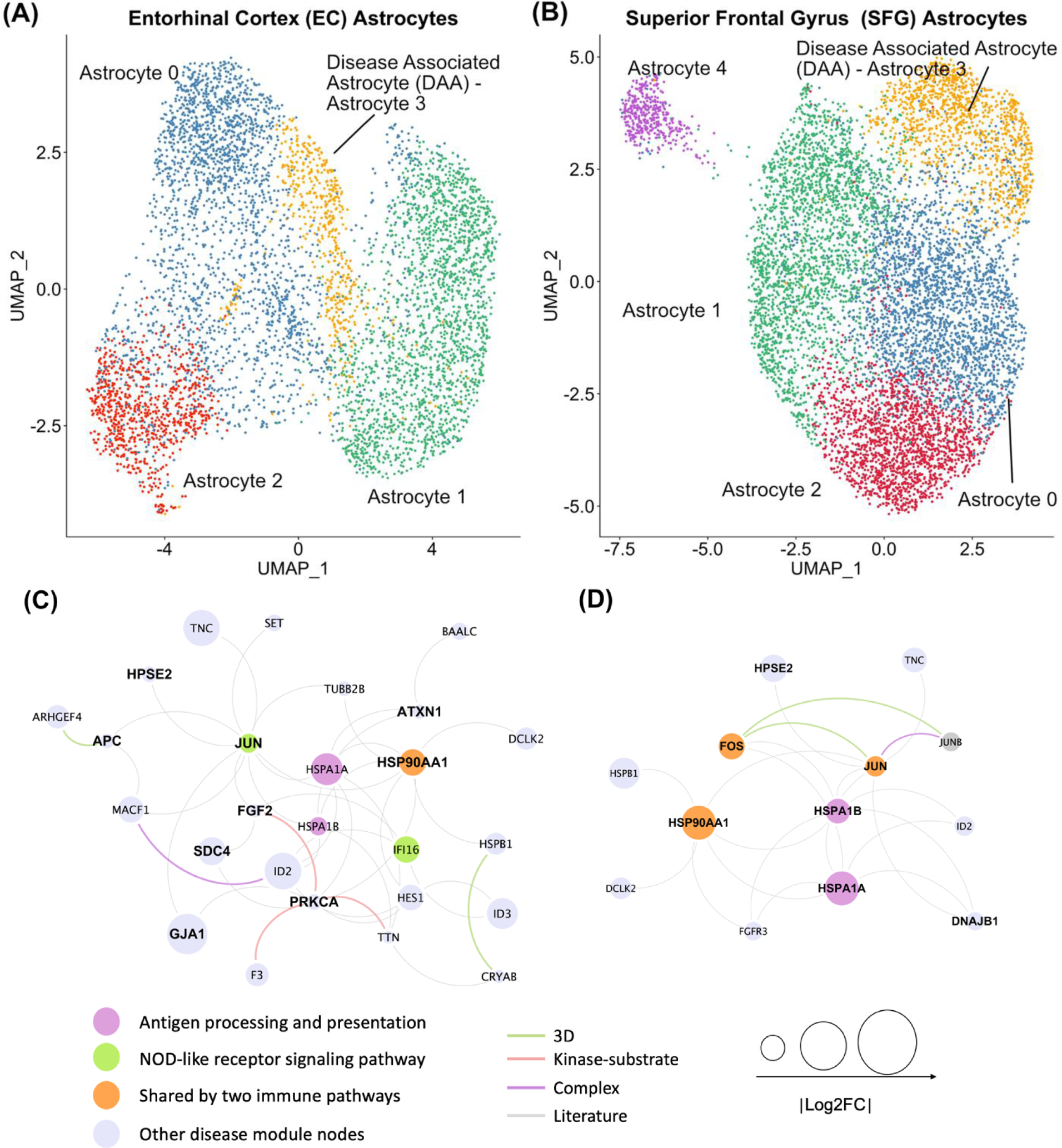
Discovery of disease-associated astrocyte (DAA) specific molecular networks for human AD brains. (**A**) Uniform manifold approximation and projection (UMAP) plot for 5599 astrocyte nuclei clustering analysis between AD patients’ brain entorhinal cortex (EC) region. (**B**) UMAP plot of clustering 8348 astrocyte nuclei for AD patients’ brain superior frontal gyrus (SFG) regions. (**C**) An identified DAA specific molecular network containing 43 protein-protein interactions (PPIs) connecting 26 genes/proteins for EC. (**D**) An identified DAA specific molecular network containing 22 PPIs connecting 13 genes/proteins for SFG. Node sizes are proportional to their corresponding |log2FC|. Nodes are color coded by KEGG immune pathways involved and edges are color coded by different experimental evidences of PPIs (**Method**).

We next inspected human brain region-specific molecular networks in DAA. Molecular networks (hDAAECnet and hDAASFGnet) of two human brain regions (EC and SFG) share 8 genes: *DCLK2, HPSE2, HSP90AA1, HSPA1A, HSPA1B, HSPB1, ID2, JUN* and *TNC* (**Figure 4C** and **4D**). There are 17 genes (e.g., *APC, ATXN1, FGF2,* and *GJA1*) and 4 genes (e.g., *DNAJB1* and *FOS*) exclusively belonging to hDAAECnet and hDAASFGnet, respectively. Adenomatous polyposis coli protein (*APC*^36^), ataxin-1-like (*ATXN1*^37^) and fibroblast growth factor 2 (*FGF2*^38^) alter AD pathogenesis by regulating beta-secretase 1 (*BACE1*), an enzyme responsible for A*β* deposition. Gap junction alpha-1 protein (*GJA1*) was reported as an AD regulator by checking 29 transcriptomic and proteomic datasets from post-mortem AD and normal control brains^39^. DnaJ homolog subfamily B member 1 (*DNAJB1*) was reported to be involved in protein folding abnormalities relevant to AD pathogenesis^40^.

We next turned to perform functional pathway enrichment analysis. As expected, we found that genes identified in DAAnets were significantly enriched in multiple key immune pathways (**Supplementary Tables 3** and **4**). To be specific, we found that majority of them had experimental evidences (such as *NFKB1, MAPK10, FOS,* and *JUN*) of roles in regulating AD pathogenesis. We next investigated selected genes in mDAAnet, hDAAECnet and hDAASFGnet using 3 immune pathways as examples: IL-17 signaling pathway, leukocyte transendothelial migration, and antigen processing and presentation (**Supplementary Tables 3** and **4**).

#### IL-17 signaling pathway

In total, we identified 8 genes (including *NFKB1, NFKBIA, MAPK10, FOS,* and *JUN*) in the IL-17 signaling pathway enriched by either mDAAnet or hDAASFGnet (**Supplementary Tables 3 and 4**). Nuclear factor NF-kappa-B p105 subunit (*NFKB1*) and NF-kappa-B inhibitor alpha (*NFKBIA*) are two regulators of the NF *κ* B signaling pathway regulating transcription of cytokines and chemokines in astrocytes. These pro-inflammatory molecules can further result in cellular damage or accelerate the production of Aβ in astrocytes^41^. The c-Jun N-terminal kinase 3 (*JNK3*), also known mitogen-activated protein kinase 10 (*MAPK10*), stimulates A*β* production and potentiates formation of neurofibrillary tangles, comprising a target for AD treatment^42^. Proto-oncogene c-Fos (*FOS*) and transcription factor AP-1 (*JUN*) are transcriptional factors regulating expression of multiple genes. Enhanced immunoreactivities of *JUN* and *FOS* were observed in AD brains, and their immunoreactivities were colocalized with paired helical filament-1 (PHF-1) within neurons, suggesting functional roles of *JUN* and *FOS* in AD pathobiology^43^.

#### Leukocyte transendothelial migration

We identify 11 mDAAnet-genes (such as *ICAM1, VCAM1, FAS, PIK3R1* and *TNFRSF1A*) in the leukocyte transendothelial migration pathway (**Supplementary Table 3**). Both intercellular adhesion molecule 1 (*ICAM1*) and vascular cell adhesion protein 1 (*VCAM1*) expression was reported to be increased by Aβ^44^. Evidence suggests that *ICAM1* and *VCAM1* facilitate leukocyte transendothelial migration, initiate endothelial signaling and affect neuroinflammation, supporting their possible roles in AD therapeutic discovery^44^. Tumor necrosis factor receptor superfamily member 6 (*FAS*) plays multiple roles in AD, including involvement in apoptosis^45^ and inflammatory processes^46^. Phosphatidylinositol 3-kinase regulatory subunit alpha (*PIK3R1*) is associated with AD and dysfunction of the insulin signaling pathway^47^. Tumor necrosis factor receptor superfamily member 1A (*TNFRSF1A*) was supported as an AD risk factor by a genome-wide haplotype-based association study in Caribbean Hispanic individuals^48^.

#### Antigen processing and presentation

We computationaly identified 7 genes (*HPSE2, FGF2, SDC4, PRKCA, HSP90AA1, HSPA1A,* and *HSPA1B*) in the antigen processing and presentation pathway enriched in either hDAAECnet or hDAASFGnet (**Supplementary Table 4**). Heat shock protein HSP 90-alpha^32^ (*HSP90AA1*) and *FGF2*^38^ have been identified as involved in AD biology. Inhibition of protein kinase C (PKC*α*) prevents A*β* from impairing synaptic activity in hippocampus in mouse model^49^. Syndecan-4 (*SDC4*), together with syndecan-3 (*SDC3*), were found to trigger fibrillation of *Aβ*1-42 in amyloid plaques^50^. Inactive heparanase-2 (*HPSE2*) was found to be over-expressed in AD human brains by a stage-dependent form^51^. Inhibiting *HPSE2* activates *HPSE* to decrease neurotoxicity and reduce tau hyperphosphorylation in AD^51^. Both heat shock 70 kDa protein 1A (*HSPA1A*^52^) and heat shock 70 kDa protein 1B (*HSPA1B*^53^) regulated oxide stress in either mouse model or human AD brains, suggesting their crucial role in AD biology and possible treatment approaches.

Two immune pathways, antigen processing and presentation and NOD-like receptor signaling, are enriched in both human brain region-specific molecular networks (hDAAECnet and hDAASFGnet, **Supplementary Table 4**). Altogether, DAA-specific molecular networks identified here are significantly enriched by known AD-associated genes and immune pathways. These DAA specific networks offer molecular mechanisms underlying AD pathogenesis and potential drug targets for treatment development.

### Alzheimer’s conserved molecular networks between microglia and astrocytes

We next compared the molecular networks between DAM and DAA (**Figure 5A**) to illustrate the unique and common disease relevant biology for microglia and astrocytes. To quantify the network relationship between DAM and DAA in the human protein-protein interactome, we use a network proximity measurement described in our recent study^54^. A higher network proximity (quantified by a lower z-score) represents a strong network relationship. We found that the closer network proximities between DAA and DAM in the human interactome compared to two random constructed networks with the same degree distributions across different network-based measurements (**Supplementary Table 5**). For instance, using the shortest path-based network proximity^54^ (**Methods**), we found a statistically significant network proximity between DAM and DAA: 1) scDAMnet and mDAAnet (z-score = -3.47, p < 1x10^-6^ [permutation test]) and 2) snDAMnet and mDAAnet (z-score = -3.31, p < 1x10^-6^ [permutation test]). These network observations indicate a strong molecular network relationship between microglia and astrocyte, which is warranted to be tested experimentally in the future.

**Figure 5.**
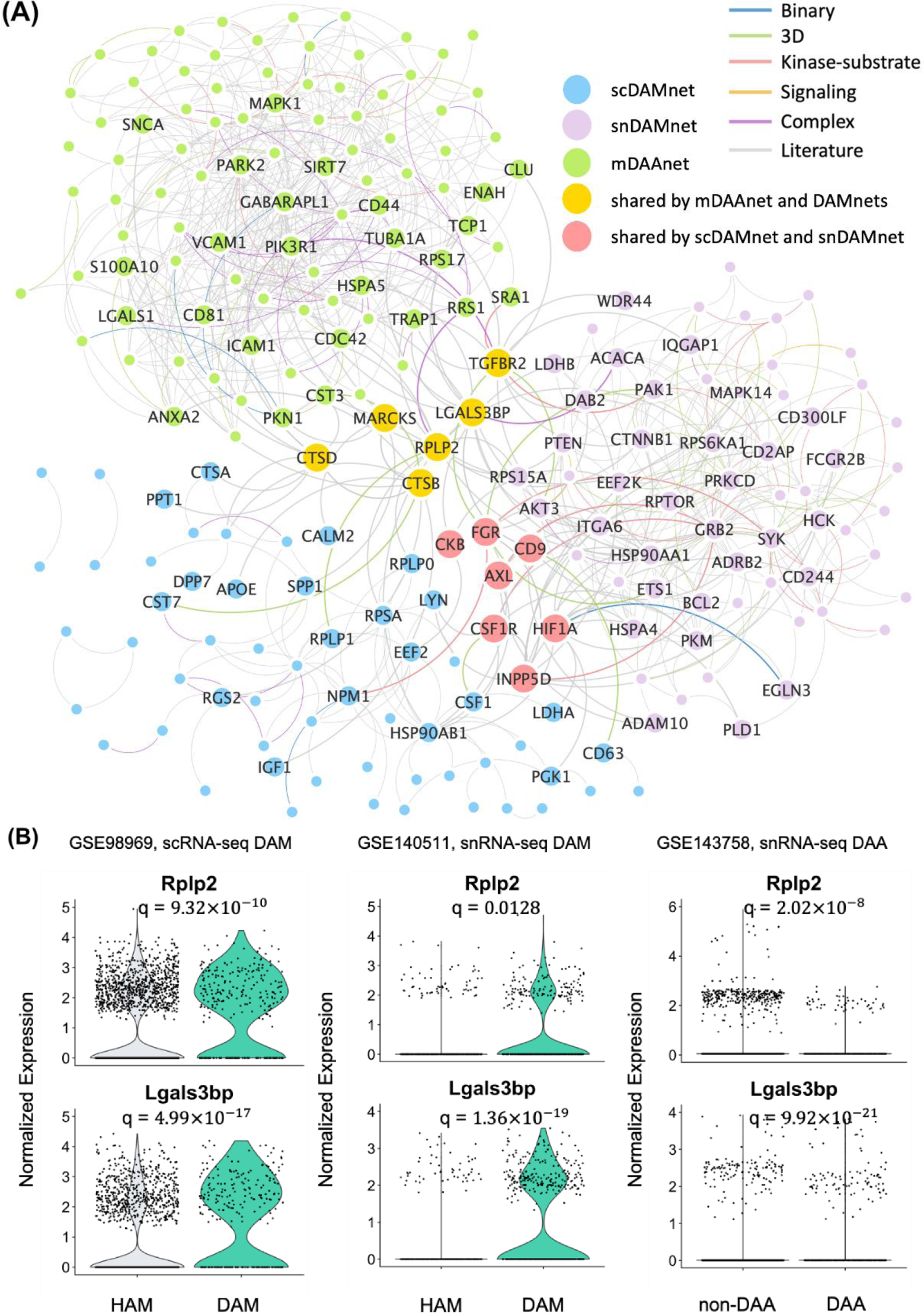
Comparison of molecular networks between disease-associated astrocytes (DAA) and microglia (DAM). (**A**) Visualization of interplays between DAM and DAA molecular networks in the human protein-protein interactome network model. (**B**) Expression levels of *LGALS3BP* and *RPLP2* for homeostatic associated microglia (HAM) versus DAM and DAA versus non-DAA. The FDR (q value) is computed using the Seurat R package (**Methods**). All details for gene differential expression analyses are provided in **Supplementary Tables 1-3**.

To be specific, we found 5 overlapped genes (*CTSB*, *CTSD*, *LGALS3BP*, *MARCKS* and *RPLP2*) and 9 commonly enriched immune pathways for molecular networks between DAM and DAA: Fc gamma R-mediated phagocytosis, B cell receptor signaling pathway, T cell receptor signaling pathway, Fc epsilon RI signaling pathway, C-type lectin receptor signaling pathway, chemokine signaling pathway, Th17 cell differentiation, leukocyte transendothelial migration, and NOD-like receptor signaling pathway. Two immune pathways (Fc gamma R-mediated phagocytosis and chemokine signaling pathway) are enriched in both scDAMnet and mDAAnet. Except *LGALS3BP* and *RPLP2* (**Figure 5B**), another 7 genes (*AXL, CD9, CKB, CSF1R, FGR, HIF1A* and *INPP5D*) are shared between scDAMnet and snDAMnet (**Figure 5A**). Three immune pathways – natural killer cell mediated cytotoxicity, platelet activation, and hematopoietic cell lineage are uniquely enriched in snDAMnet, while IL-17 signaling pathway, Toll-like receptor signaling pathway, Th1 and Th2 cell differentiation, and RIG-I-like receptor signaling pathway are exclusively enriched in mDAAnet (**Table 2**). In summary, microglia and astrocyte may synergistically trigger neuroinflammation in AD in a cell type-specific manner.

### Metabolites trigger molecular networks between astrocytes and microglia

Since AD is a pervasive metabolic disorder that is linked with altered immune responses^55^. We inspected the network relationship (**Methods**) between well-known metabolic genes collected from Kyoto Encyclopedia of Genes and Genomes^56^ (KEGG, **Methods**) and the identified molecular networks from both DAM and DAA. We found that metabolic genes have a closer network relationship with DAM- or DAA-molecular networks compared to randomly selected genes after adjusted degree (connectivity) bias in the human interactome (**Methods** and **Supplementary Table 5**). We therefore turned to investigate whether environmental factors (including metabolites) trigger molecular network perturbation between astrocytes and microglia. We performed an integrative network-based analysis of AD-related metabolite-enzyme associations and the human protein-protein interactome. We constructed a heterogenous networks, including 373,320 edges consisting of 26,990 metabolite-enzyme associations and 346,330 PPIs (**Methods**). Specifically, we assembled 155 AD-related metabolites (**Supplementary Table 6**) supported by experimental evidence and found in human brain, blood and cerebrospinal fluid samples from 12 well-performed clinical studies (**Supplementary Table 6**) as well as a high-quality database of small molecule metabolites, the Human Metabolome Database^57^ (HMDB) (**Methods**). Based on our observations, we performed graphic computations on the heterogenous network and extracted a subgraph consisting of 251 nodes and 1,404 edges as the DAM-DAA networks (**Methods, Figure 6A** and **Supplementary Figure 7A**).

**Figure 6.**
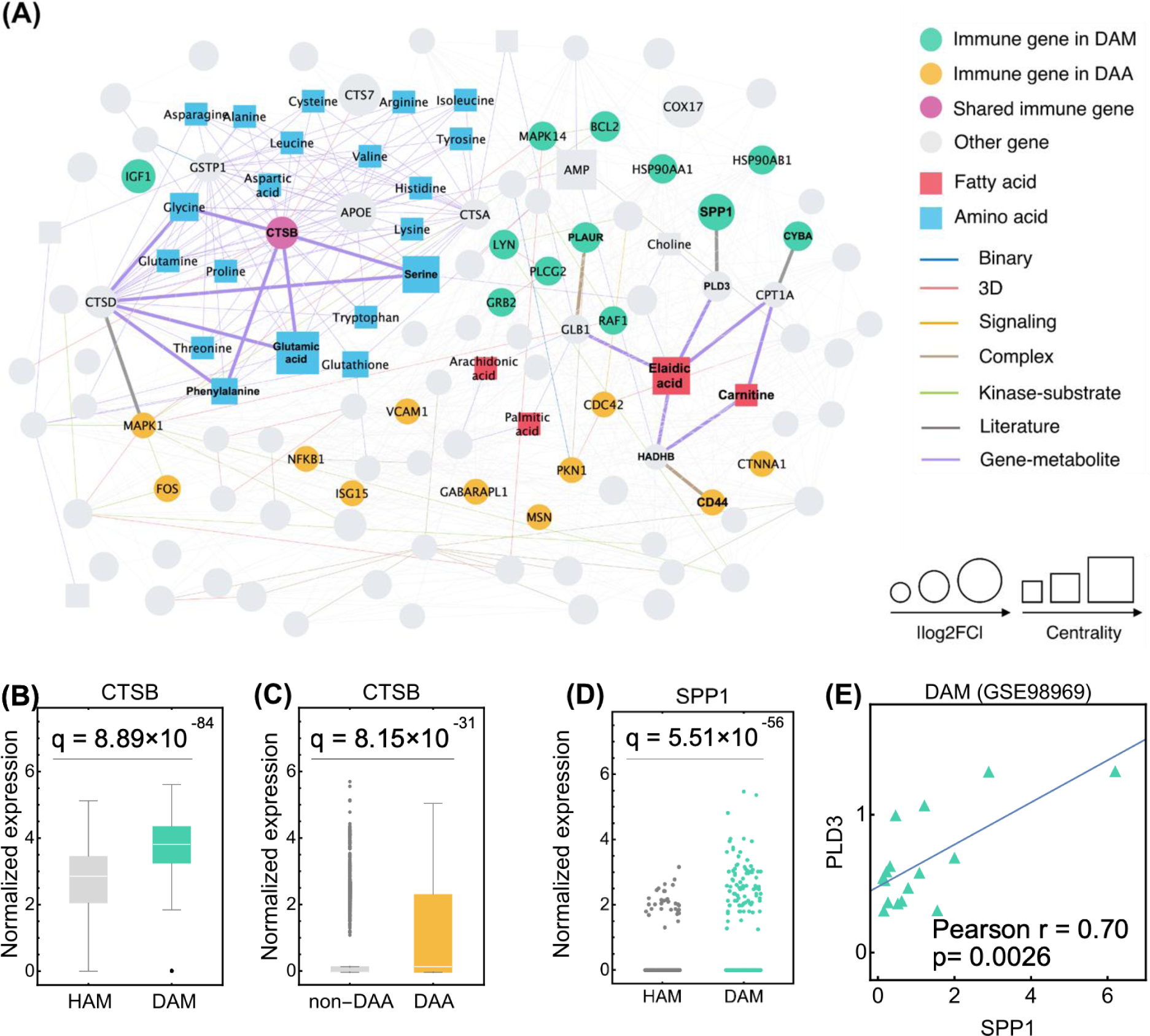
A metabolite-triggered molecular network between disease-associated astrocytes (DAA) and microglia (DAM). (**A**) A highlighted subnetwork of the metabolite-enzyme network between DAM and DAA in the human protein-protein interactome network model; (**B** and **C**) Expression of *CTSB* is significantly elevated in (**C**) DAM and (**D**) DAA, compared to homeostatic associated microglia (HAM) and non-DAA, respectively. (**D**) Expression of *SPP1* is significantly elevated in DAM compared with HAM. Each dot represents one cell; (**E**) Pearson correlation analysis shows that *SPP1* and *PLD3* have the coordinated change trends in DAM. Gene expression is counted by the average Unique Molecular Identifier (UMI) count.

In total, we found 70 enzymes that regulated the AD-related metabolites: i) 50 (e.g. *APOE*, *GLB1*, *LDHB* and *PLCG2*) enzyme-coding genes from DAM; ii) 23 (e.g. *SIRT7*, *HADHB*, *MAPK1* and *PGAM1*) genes from DAA, and iii) 3 (*CTSB*, *CTSD* and *TGFBR2*) genes which are common to both DAM and DAA (**Supplementary Figure 7B** and **Supplementary Table 6**). For instance, the *CTSB*, encoding cathepsin B, involved in catabolism and immune resistance in humans^58^, has elevated expression (**Figures 6B** and 6**C**) in both DAM (Fold-Change [FC] = 2.48, q = 8.89×10^-84^) and DAA (FC = 1.84, q = 8.15×10^-31^). The pathway enrichment analysis on these 70 enzymes shows the effect of AD on metabolic homeostasis (e.g., glycolysis and gluconeogenesis) and highlights some immune signaling pathways such as IL-3 and IL-5 (**Supplementary Figure 7C**). These observations provide the proof-of-concept of metabolism-driven immune responses between microglia and astrocytes in AD pathogenesis.

Using a betweenness centrality measure (**Methods, Supplementary Table 6**), we found that fatty acids and amino acids (**Figure 6A**) were two primary types of metabolites involved in molecular networks between the DAM and DAA. For example, SPP1^59^ and CD44^60^, two cellular molecules that promote chronic inflammatory diseases, are significantly over-expressed in both DAM (FC = 5.35, q = 5.51×10^-56^, **Figures 6D**) and DAA (FC = 1.30, q = 5.13×10^-12^) compared to HAM and non-DAA, respectively. Elaidic acid, a major trans-fat, shows the largest centrality among all metabolites and is connected with SPP1 and CD44 by two enzymes involved in fatty acid metabolism, including phospholipase D (PLD3) and hydroxyacyl-CoA dehydrogenase (HADHB), respectively^61^. Co-expression analysis uncovers the coordinated change trends of SPP1 and PLD3 in DAM (Pearson r = 0.70, *P* = 0.0026, **Figure 6E**). Meanwhile, carnitine, transporting the long-chain fatty acids into mitochondria for oxidation, is also involved in the interactions with differentially expressed genes in both DAA (i.e., CD44) and DAM (such as *CYBA* and *PLAUR*). These findings suggest the potential bridge roles of fatty acid metabolism for the communication between astrocytes and microglia under AD pathology. Moreover, amino acids, especially glutamate, serine, and phenylalanine, may trigger the immune responses by directly targeting the shared gene *CTSB* which is associated with the amyloid precursor protein processing in both DAM^8^ and DAA^10^. In summary, characterizing the network-based relationship between the DAM-/DAA-specific molecular networks and the AD-related metabolites within the human interactome network model identifies underlying immunometabolism mechanisms related to the immune interplay of astrocytes and microglia in AD triggered by cellular metabolism.

### Network-based discovery of drug candidates via reversing gene expression of microglia and astrocytes

We next turned to identify potential drug candidates by specifically targeting molecular networks in microglia and astrocytes. As shown in **Figure 1**, we collected drug-gene signatures in human cell lines from the Connectivity Map (CMap) database^62^. We posited that if a drug significantly reverses dysregulated gene expression (measured by the most up-regulated and down-regulated genes, **Methods**) of DAM or DAA involving in AD, this drug may have potential in treating AD. We performed gene set enrichment analysis (GSEA) and calculated enrichment score (ES) using permutation tests (**Methods**). We used ES > 0 and p ≤ 0.05 as the valid cutoffs to prioritize potential drug candidates. In total, we investigated 1309 drugs with known target information from the DrugBank^63^ database or having gene signatures from the CMap^62^. We obtained 172, 234, 187, 124, and 195 candidate drugs (ES > 0 and p ≤ 0.05) based on GSEA analyses using snDAMnet, scDAMnet, mDAAnet, hDAAECnet and hDAASFGnet, respectively. The complete drug prediction results are summarized in **Supplementary Table 7**. A Venn diagram showing the relations of predicted drugs among different molecular networks is presented in **Supplementary Figures 8A** and **8B**. We found that drugs with ES >0 predicted by at least one molecular network have potentially beneficial for AD across several pharmacological categories: anti-inflammatory agents, immunosuppressive agents, adrenergic beta-2 receptor agonists, adrenergic alpha-antagonists, and antipsychotic agents (**Figure 7A**). We next focused on 4 high-confidence drug candidates (**Figure 7B**) using subject matter expertise based on a combination of factors: (i) strength of the predicted associations (ES value); (ii) novelty of the predicted associations with established mechanisms-of-action (such as anti-inflammatory); (iii) literature-based evidence in support of prediction; (iv) availability of sufficient patient data for meaningful evaluation (exclusion of infrequently used medications).

**Figure 7.**
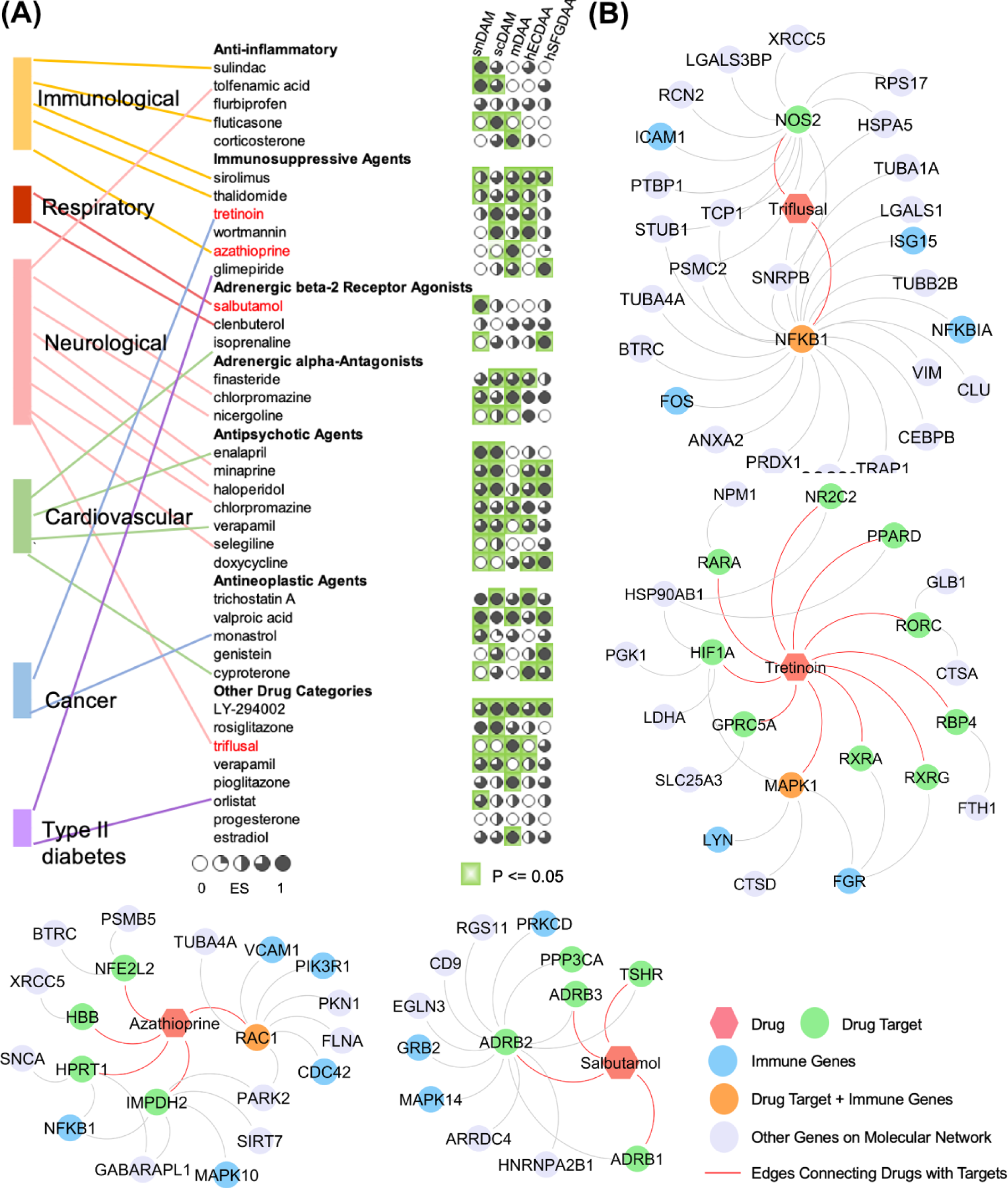
Network-based discovery of repurposable drug candidates for AD by specifically reversing gene expressions of disease-associated microglia (DAM) and disease-associated astrocytes (DAA). (**A**) Selected drugs that specifically target five different DAM or DAA molecular networks. Drug are grouped by five different classes: immunological, respiratory, neurological, cardiovascular, cancer, and type II diabetes. Four high-confidence drugs were highlighted by red text. (B) Proposed mechanism-of-actions for 4 selected drugs by drug-target network analysis.

**Figure 8.**
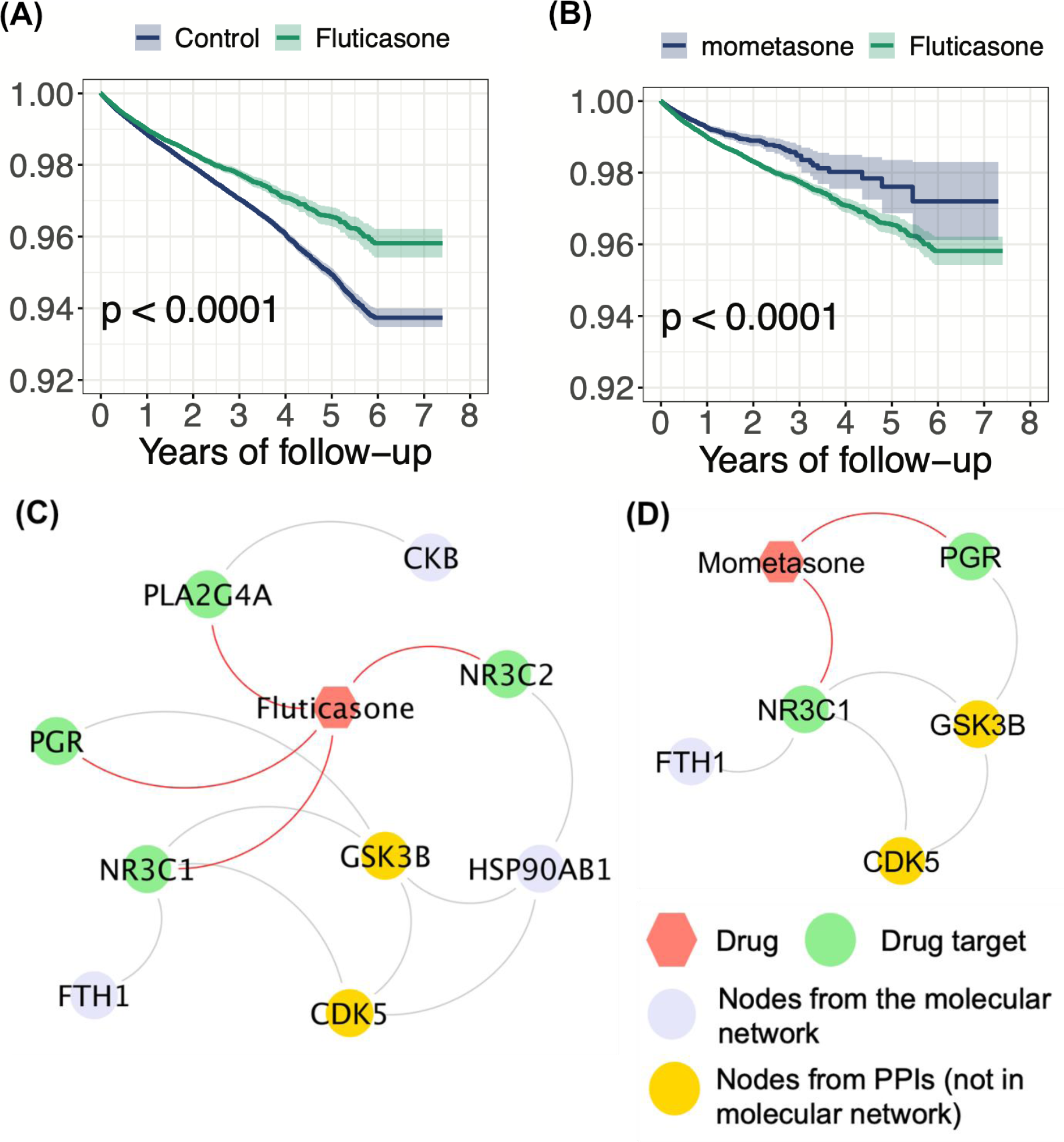
Longitudinal analysis reveals that fluticasone and mometasone reduce risk of AD incidence in patient data. Two comparison analyses were conducted including: (**A**) fluticasone vs. a matched control population and (**B**) fluticasone vs. mometasone. We estimated the un-stratified Kaplan-Meier curves, conducted propensity score stratified (n strata = 10) rank test and Cox models after adjusted all possible confounding factors, including age, gender, race, disease comorbidities (see Methods). (**C** and **D**) Proposed mechanism-of-action for treatment of AD by fluticasone and mometasone using drug-target network analysis.

#### Immunosuppressant

The observations suggested that an immunosuppressive drug (Rapmaycin) could potentially benefit AD treatment in both cellular experiments and animal models^64^. The reported underlying mechanism included regulating autophagy and cellular signaling pathways^64^. One of our top-predicted drugs – azathioprine (ranking 1^st^ from mDAAnet after removing drugs with no targets according to the DrugBank database, p = 0.02, **Supplementary Table 7**) is an immunosuppressive drug that used to treat autoimmune disorder disease - rheumatoid arthritis^65^ and prevents renal transplant rejection^66^. A study reported therapeutic potential of azathioprine in AD^67^. When integrating drug targets of azathioprine into the predicted molecular network of mDAAnet, we found that azathioprine directly targets ras-related protein Rac1 (*RAC1*), a key immune gene in astrocytes (**Figure 7B**). In addition, azathioprine indirectly targets several key immune genes, including *VCAM1, PIK3R1, CDC42, MAPK10,* and *NFKB1* in mDAAnet (**Figure 7B**). In addition to *VCAM1, MAPK10, NFKB1* and *PIK3R1* as critical AD pathological modulators, CDC42 small effector protein 1 (*CDC42*) was shown to be biologically associated with AD since it presented a large overlap with cytokine abnormalities^68^. These observations suggest that azathioprine is a candidate immunosuppressant agent by specifically targeting molecular networks in DAA.

#### Antithrombotic anticoagulan

In one AD transgenic mouse model, it was found that long term anticoagulation with dabigatran (an approved thrombin inhibitor), could help to preserve cognition, cerebral perfusion, and blood brain barrier (BBB) function and alleviate A*β* deposition^69^. Our predicted antithrombotic anticoagulant, triflusal (ranking 6^th^ from mDAAnet after removing drugs with no targets according to the DrugBank database, p < 1x10^-6^, **Supplementary Table 7**), presented a reduced risk of AD’s dementia progression in a randomized, double-blind, placebo-controlled trial with 257 subjects^70^. Another study reported that triflusal repaired defects in axonal curvature and cognition in an AD transgenic mouse model^71^. In drug-target network analyses (**Figure 7B**), triflusal directly targets *NFKB1* and indirectly targets other 4 immune genes (*ISG15, NFKB1A, FOS*, and *ICAM1)* in mDAAnet, suggesting a possible anti-inflammatory mechanism-of-actions for AD.

#### Beta2-adrenergic receptor agonist

Longitudinal and cross-sectional epidemiological studies revealed that treatment with beta-blockers reduces AD incidence in individuals suffering from hypertension^72^. From cellular experiments and animal models, it was found that β-adrenergic receptors play a role in AD pathogenesis via influencing A *β* production and inflammation^72^. Another study demonstrated in an APP/PS1 transgenic AD mouse model that beta2-adrenergic receptor activation could enhance neurogenesis and repair cognitive deficits^73^. Salbutamol (ranking 4^th^ from snDAMnet after removing drugs with no targets according to the DrugBank database, p < 1x10^-6^, **Supplementary Table 7**), a selective beta2-adrenergic receptor agonist used in the treatment of asthma, is a highly predicted candidate from our network-based approach. Salbutamol inhibits tau accumulation in vitro^74^. When incorporating its drug target with molecular network (snDAMnet), salbutamol interacts directly with 3 immune genes (*PRKCD*, *GRB2* and *MAPK14*) as shown in **Figure 7B**. Except *PRKCD* and *MAPK14* aforementioned, literature evidence also demonstrates that growth factor receptor-bound protein 2 (*GRB2*), involving in the C-terminal fragments (CTF) – ShcA complexes, plays a role in influencing AD development^75^.

#### Retinoic acid

Retinoic acid, another potential AD therapeutic option, has been widely studied^76^. One of the explanations elucidating its possible effectiveness for AD treatment is its inhibition of oxidative stress and abnormal differentiation of neurons, two common pathological factors for AD^76^. In the AD APP/PS1 transgenic mouse model, treatment with all*-trans* retinoic acid (ATRA) significantly decreased APP phosphorylation and processing, reduced microglia and astrocyte activities, down-regulated cyclin-dependent kinase 5 (*CDK5*) activity, and enhanced cognitive capabilities^77^. Tretinoin (ranking 1^st^ from scDAMnet after removing drugs with no targets according to the DrugBank database, p < 1x10^-6^, **Supplementary Table 7**), a US Food and Drug Adminisatrion (FDA)-approved drug for acute promyelocytic leukemia (APL), is one of our highest predictions^78^. Experiments with AD transgenic mouse models showed that tretinoin decreased activation of microglia and astrocytes^77^. Mechanistically, tretinoin directly targets mitogen-activated protein kinase 1 (*MAPK1*), *LYN* and *FGR* in the scDAMnet (**Figure 7B**).

### Validating possible causal associations in patient data

Fluticasone, a synthetic glucocorticoid which is FDA-approved for several inflammatory indications, is one of our top predicted drugs based on scDAMnet. We evaluated the fluticasone user’s vulnerability to AD by analyzing 7.23 million U.S. commercially insured individuals (MarketScan Medicare supplemental database, see Methods). We conducted two cohort analyses to evaluate the predicted association based on individual level longitudinal patient data and state-of-the-art pharmacoepidemiologic methods. These included: (i) fluticasone vs. a matched control population (non-fluticasone user), and (ii) fluticasone vs. mometasone (an FDA-approved corticosteroid for skin conditions, hay fever, and asthma). For each comparison, we estimated the un-stratified Kaplan-Meier curves and conducted propensity score stratified (n strata = 10) log-rank tests and the Cox regression model.

We found that individuals taking fluticasone were at significantly decreased risk for development of AD (hazard ratio (HR) = 0.858, 95% confidence interval [CI] 0.829-0.888, *P* < 0.0001, **Figure 8A**) in a retrospective case-control validation. Importantly, propensity score matching cohort studies confirmed mometasone’s association with reduced risk of AD in comparison to fluticasone (HR =0.921, 95% CI 0.862-0.984, *P* < 0.0001, **Figure 8B**). Another independent database – FDA MedWatch Adverse Events Database revealed that the combination of fluticasone and ibuprofen could be a therapeutic option for AD^79^. To infer the potential mechanisms-of-action of fluticasone in AD, we integrate drug target network, the extracted molecular network and human PPIs (scDAMnet, **Figure 8C**). Network analysis shows that fluticasone could indirectly target GSK3B and CDK5 (**Figure 8C**). And literature study demonstrates that GSK3B and CDK5 are two most relevant targets for AD^80^.

## Discussion

The emergence of single-cell/nucleus sequencing technologies and development of computational tools enables us to capture new insights into neuroinflammation for AD from the molecular network perspective. In this study, we systematically reconstructed molecular networks for both DAM and DAM by uniquely integrating scRNA/sn-RNA-seq profiles form both AD transgenic mouse models and human AD brains. We showed that in AD, affected genes regulate either one (such as *CSF1R, CD33, CCL3/4* and *CCR5*) or multiple (e.g., *SYK, ICAM1, VCAM1, NFKB1, HSP90AA1, JUN* and *FOS*) immune pathways. The enriched immune pathways and network proximity^54^ analyses indicate that microglia and astrocytes may share a strong network relationship in the human protein-protein interactome (**Figure 5A** and **Supplementary Table 5**). Via incorporating the enzyme-metabolite associations, we found that fatty acids (e.g., elaidic acid) and amino acids (e.g., glutamate, serine and phenylalanine) may trigger immune alterations between DAM and DAA. Finally, we computationally identified that existing drugs (including azathioprine, triflusal, salbutamol, and retinoic acid) offer potential candidates for AD by reversing gene expression of DAM or DAA. Importantly, we demonstrated that fluticasone and mometasone were significantly associated with the decreased risk of AD in a large-scale patient data.

We acknowledged several potential limitations in the current study. Although two snRNA-seq and scRNA-seq datasets of DAM present consistent gene expression patterns (**Supplementary Tables 1 and 2**), snDAMnet and scDAMnet showed a small gene overlap. There are several possible explanations. For example, single-cell and single-nucleus may generate different cell abundances during cell processing. DAM accounts for around 12% of microglia based on scRNA-seq data in 5XFAD mice, while percentage surges to more than 50% based on snRNA-seq data in 5XFAD mice (**Tables 1A** and **1B**). For DAA, both mouse (**Supplementary Table 3**) and human (**Supplementary Table S4**) RNA-sequencing data display partial consistent gene expression patterns, including DAA marker genes *GFAP*, *CD44*, *HSPB1*, *APOE* and *TREM2*. Several opposite human marker genes’ expression patterns are also detected when compared with mouse data, such as *TNC*, *SLC1A2*, *SLC1A3*, and *GLUL*. Two human molecular networks (hDAAECnet and hDAASFGnet) built from two human brain regions are similar. The network proximity analyses under different measurements also show that the distances between two human molecular networks are significantly closer compared to distances between random networks with specific restrictions (**Methods** and **Supplementary Table 5**). However, when comparing between the mouse and human molecular networks, their overlaps are very small. Samples collecting from different brain regions is one reasonable explanation. The human RNA-sequencing data we used were collected from entorhinal cortex and superior frontal gyrus; while the mouse data were collected from hippocampus.

Network proximity evaluations show that the distances between human and mouse molecular networks are small; however, less significant when compared to distances among mouse molecular networks (**Supplementary Table 5**). One study showed that immunology in human AD brains and mouse models were different^81^. Another more recent study argued that gene signatures were very distinct between mouse model 5XFAD and human AD brains in DAM as well^8^. For example, upregulated 5XFAD DAM marker genes, Lpl and Cst7, could not be detected in human AD brain^82^.

These unresolved must be addressed in the future. Quality of AD outcomes or phenotypes defined by current clinical criteria (such as Braak score) may influence our findings as well. According to the National Institute of Neurological Disorders and Stroke-Alzheimer’s Disease and Related Disorders Association (NINDS-ADRDA), Braak stage 2 is probable AD with supported evidence, and Braak stage 6 is considered as definite AD. For two brain regions, EC and SFG, there are no apparent differences of normalized nucleus abundance percentage across different Braak stages for both DAA and non-DAA (**Tables 1** and **Supplementary Figure 2D** and **2E**). Finally, incompleteness and potential biases of human protein-protein interactcome and drug-target networks may influence our network-based findings as well.

In summary, we proposed a network-based approach that incorporates snRNA-seq and scRNA-seq data sampled from either mouse models or AD patient brains, PPIs, enzyme-metabolite associations, and drug target networks, along with the large-scale patient validation database. We showed the molecular networks derived from DAM and DAA are significantly enriched for various well-known immune pathways and AD pathobiological pathways. We showed that the identified molecular network from DAM and DAA offer potential targets for drug repurposing, which we validated for proof-of-concept in a large-scale, real-world patient database. In summary, we believe that the network-based approach presented here, if broadly applied, would significantly catalyze innovation in AD drug discovery and development for AD and other disease by utilizing the large-scale existing omics data at the single-cell/nucleus levels.

## Methods and Materials

### Single-cell and nucleus RNA-sequencing

We collected single cell and nucleus transcriptomics data from four recently published papers. The complete mouse single-cell transcriptomics data were sequenced from various transgenic mouse models, including C57BL/6, 5XFAD, Trem2 knock out C57BL/6, Trem2 knock out 5XFAD and SOD1 and different organs, i.e., whole brain, cortex, cerebellum and spinal cord with different ages, i.e., 7 weeks, 80 and 135 days, 1, 3, 6, 9 and 20 months. In this study, we utilized data from 16 C57BL/6 (whole brain) and 16 5XFAD 6 month-mouse. In total, there were 12,288 cells sequenced from 32 mouse samples. Two of three snRNA-seq data were collected from mouse samples as well (GSE140511 and GSE143758). Dataset GSE140511 contained four transgenic male mouse models, including C57BL/6, 5XFAD, Trem2 knock out C57BL/6 and Trem2 knock out 5XFAD. In this study, we considered the 7-month mouse models which in total sequenced 90,647 nuclei as described in the original literature^82^. The second mouse nucleus dataset GSE143758 contains two transgenic mouse models C57BL/6 and 5XFAD from both hippocampus and cortex regions and with different ages. Similarly, we utilized in total 50,242 nuclei data from the 7-month mouse models with 5 5XFAD and 6 C57BL/6 samples^10^. Finally, the human single-nucleus transcriptomics data^83^ contains ten male frozen post-mortem human brain tissues from both superior frontal gyrus and entorhinal cortex regions. There are 3, 4 and 3 human brains tissues with Braak stages 0, 2, and 6, respectively. The raw data (including astrocytes, excitatory neurons, inhibitory neurons and microglia cells) downloaded from on Gene Expression Omnibus (GSE147528). All the following data analysis are based on the processed data after quality control, and there are 5599 and 8348 nuclei for samples from entorhinal cortex and superior frontal gyrus, separately.

### Bioinformatics analysis of single cell/nucleus RNA-sequencing data

The analyses were completed with Seurat^84^ (v3.1.5), scran^85^ (v1.16.0), scater^86^ (v1.16.1) packages in R with steps complied with the original literatures. Data were normalized using a scaling factor of 10,000 and all DEG analyses are conducted by function *FindMarkers* in Seurat^84^ R package with parameter *test.use = ‘MAST’*. The detailed data analysis steps for each dataset (GSE98969, GSE140511, GSE143758 and GSE147528) are illustrated in sequences.

**GSE98969** The data used are from whole brain cells of 6 months 5XFAD (16) and C57BL/6 (16) mice that express gene CD45. For quality control, cells with mitochondrial content >5% and UMIs < 500 were removed. Genes with mean expression smaller than 0.005 UMIs/cell were discarded for analysis. Data were normalized using a scaling factor of 10,000 and functions *FindIntegrationAnchors* and *IntegrateData* in Seurat^84^ R package are used for batch effect correction for samples collected from different plates. Principle component analysis was performed using the top 2000 most variable genes and clustering was performed using the top 30 principal components (PCs) and resolution of 0.4. After identifying clusters for DAM and HAM, separately (gene markers, see **Figure 1b**^8^), DEGs are compared between DAM and HAM by considering cells from 5XFAD mice only. The whole pipeline was completed with Seurat^84^ R package.

**GSE140511.** The process for clustering different cell types are provided in the original literature^82^. We used microglia nuclei and reproduce the clustering procedures to isolates DAM and HAM nuclei. Considering all microglia nuclei, principle component (PC) analysis was performed using the top 3000 most variable genes and sub-clustering was performed using the top 10 PCs and resolution of 0.1. Again, after identifying clusters enriched in DAM and HAM nuclei (gene markers, **Figure 1b** in^8^), DEGs are compared between DAM and HAM by considering nuclei from 5XFAD mice only. The whole pipeline was completed with Seurat^84^ R package.

**GSE143758.** The process for clustering different cell types are provided in the original literature^10^. In this study, we used astrocyte nuclei and reproduced the clustering procedures to isolate DAA) and non-DAA nuclei. Considering all astrocyte nuclei, principle component analysis was performed using the top 2000 most variable genes and sub-clustering was performed using the top 10 PCs and resolution of 0.3. After identifying clusters enriched in DAA nuclei by comparing the expression pattern of marker genes (**Figure 1e** in^10^) among sub-clusters. We computed DEGs between DAA and non-DAA by considering nuclei from 5XFAD mice. All analyses were performed using Seurat^84^ R package.

**GSE147528.** We used astrocyte cell subtype - reactive astrocyte, which is associated with AD^83^. We considered astrocyte nuclei and clustering analysis was first performed by *quickCluster* function and size factors were computed by *computeSumFactors* function with parameter *min.mean = 0.1* in scran R package. Then count matrix was normalized by the computed size factors and log-transformed by function *logNormCounts* in scater R package. Top 1000 highly variable genes were selected by functions *modelGeneVar* and *getTopHVGs* in scran R package. Functions *FindIntegrationAnchors* and *IntegrateData* in Seurat^84^ R package were used for batch effect correction, and clustering was performed using the top 12 PCs and resolution of 0.2. After identifying clusters enriched in reactive astrocyte nuclei (gene markers^83^, **Figures 2a and 2b**), DEGs are compared between reactive astrocytes and non-reactive astrocytes for nuclei from both superior frontal gyrus and entorhinal cortex regions, respectively.

### Building Human Protein-protein interactome

To build the comprehensive human interactome from the most contemporary data available, we will assemble 18 commonly used PPI databases with experimental evidence and the in-house systematic human PPI we have previously utilized^3^: (i) binary PPIs tested by high-throughput yeast-two-hybrid (Y2H) systems^87^; (ii) kinase-substrate interactions by literature-derived low-throughput and high-throughput experiments from KinomeNetworkX^88^, Human Protein Resource Database (HPRD)^89^, PhosphoNetworks^90^, PhosphositePlus^91^, DbPTM 3.0 and Phospho.ELM^92^; (iii) signaling networks by literature-derived low-throughput experiments from the SignaLink2.0^93^; (iv) binary PPIs from three-dimensional protein structures from Instruct^94^; (v) protein complexes data (∼56,000 candidate interactions) identified by a robust affinity purification-mass spectrometry collected from BioPlex V2.0^95^; and (vi) carefully literature-curated PPIs identified by affinity purification followed by mass spectrometry from BioGRID^96^, PINA^97^, HPRD^98^, MINT^99^, IntAct^100^, and InnateDB^101^. As of December 2019, the updated human interactome constructed includes 351,444 PPIs connecting 17,706 unique proteins.

### Description of GPSnet

GPSnet algorithms takes two inputs: node score and one background PPI network. The node score was defined as follows: for DEGs with q <= 0.05, the node scores equal to absolute value of log2FC. In order to generate a module, GPSnet starts with a randomly selected gene/protein (node) as the seed gene. During each iteration, one of candidate genes (1^st^ order neighbors of current seed genes) that satisfying the following two conditions at the same time will be added: (1) a p-value of the connectivity significance P(i) (**Eq. 1**) is less than 0.01; (2) the updated module score is greater than the current one (**Eq. 2**). We repeated steps (1) and (2) until no more genes (nodes) can be added. In this study, we built ∼100,000 raw modules ranked by module scores. For each raw module, the corresponding module score can be computed (**Eq. 2**) and all raw modules are ranked in decreasing module score order and the protein frequency is defined based on truncated raw modules. We generated the final network modules by assembling top raw network modules (**Supplementary Tables 1-4**).

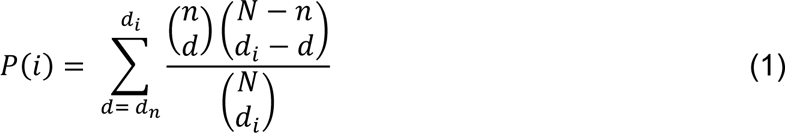

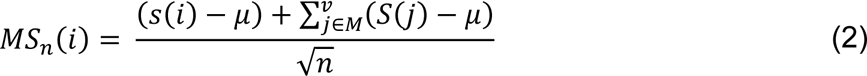

Where, N denotes all proteins/genes in the PPI, n represents numbers of nodes in the module, *d*_*n*_ is the numbers of neighbors of gene i, *d*_*i*_ is the degree of gene i, *MS*_*n*_(*i*) denotes the updated module score if adding node i, M denotes the current module, and μ is the average node score (|log2FC| in this work) of all genes with respect to the PPI.

### Network proximity

To quantify the relationships of two molecular networks (DAM vs. DAA) in the human interactome, we adopted the shortest path-based network proximity measure^54^ as below.

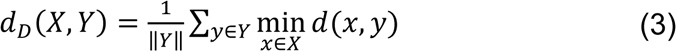

where *d*(*x*, *y*) is the shortest path length between gene *x* and *y* from gene list *σ* (DAM) and *Y* (DAA), respectively. To evaluate whether such proximity was significant, the computed network proximity is transferred into z score form as shown below:

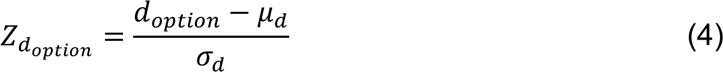

Here, *μ*_*d*_ and σ_*d*_ are the mean and standard deviation of permutation test with 1000 random experiments. In each random experiment, two random subnetworks *X*_*r*_ and *Y*_*r*_ are constructed with the same numbers of nodes and degree distribution as the given 2 subnetworks X and Y, separately, in the human protein-protein interactome.

### Network analysis metabolite-enzyme associations

We collected 136 AD-related metabolites from 12 studies and the Human Metabolome Database (HMDB)^57^. All metabolistes were identified in AD-related human samples, including brain tissue, cerebrospinal fluid, and blood. All of these results are free available in our website AlzGPS (https://alzgps.lerner.ccf.org/). We collected experimentally reported metabolite-enzyme associations from three commonly used data sources, including KEGG^56^, Recon3D^103^, and HMDB^57^, and assembled them with the human PPI network. The updated network contains 373,320 links connecting with 17,826 unique proteins (including enzymes) and 1,419 metabolites. Then we mapped 224 DAM and DAA disease module genes and the 155 AD-related metabolites to the new network and computed the maximal subgraph: (1) we found 614 unique nodes which were the first or second order neighbors of 61 DAM and DAA immune genes; (2) we obtained 71 metabolites by considering the intersection of 614 unique nodes and 155 AD-related metabolites; (3) a subnetwork connecting 224 genes and 71 metabolites was generated. Finally, we computed the network paths that connected the DAM and DAA genes using the large betweenness centrality.

### Connectivity Map (CMap) and DrugBank database

The Connectivity Map data used in this study contains 6100 expression profiles relating 1309 compounds^62^. A parameter *α* defined below is used to leverage the extent of differential expression for a given set of genes.

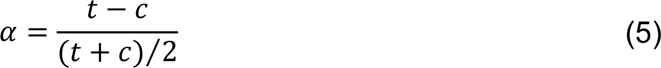

Here t is the scaled and thresholded average difference value for the drug treatment group and c is the thresholded average difference value for the control group. Therefore, a zero *α* value indicates no expression change after drug treatment, and a positive *α* value means elevated expression level after drug treatment and vice versa. Drug gene signatures with α > 0.67 (0.67 equals the 2-folds increment) are considered as up-regulated drug-gene pairs, and α < -0.67 are denoted as down-regulated drug-gene pairs.

### Gene Set Enrichment Analysis (GSEA)

We utilized GSEA algorithm to predict drugs for each cell subtype. GSEA algorithms takes two inputs: CMap database and the extracted molecular network. Detailed descriptions of GSEA have been illustrated in^104^. To be specific, the GSEA enrichment score (ES) is calculated as shown below.

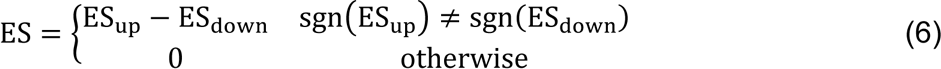

Both *ES*_*up*_ and *ES*_*down*_ are computed for up- and down-regulated genes in input molecular network separately with the same scheme as shown below in a 2-step manner. We first compute intermediate parameters *a* and *b*:

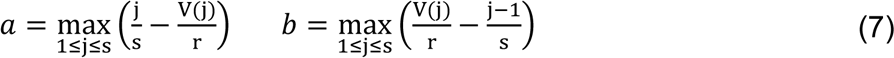

where j = 1, 2, …, s were the gene sets from molecular network sorted in ascending order by their rank in the gene profiles of the drug being evaluated. The rank of gene j is denoted by V(j), where 1 ≤ *V*(*j*) ≤ *r*, with r being the number of genes (12,849) from the drug profile. Then, the corresponding *ES*_*up*_ and *ES*_*down*_ equal:

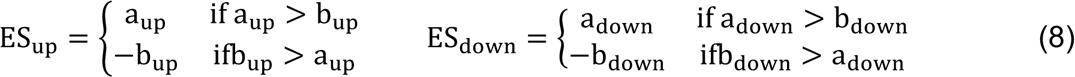

In the above equations, *a*_*up*/*down*_ and *b*_*up*/*down*_ are computed with respect to up- and down-regulated genes in molecular network, separately. The GSEA ES represents drug potential capability to reverse the expression of the input molecular network. Permutation tests repeated 100 times using randomly gene lists consisting of the same number of up- and down-regulated genes as the input molecular network were performed to leverage the significance of the computed ES value. Therefore, drugs with large positive ES value and significant p (p ≤ 0.05) were selected.

### Enrichment Analysis

All pathway and disease enrichment analyses in this study are conducted via either KEGG 2019 Mouse or KEGG 2019 Human and DisGeNET^18^ from Enrichr^105^, respectively. The DisGeNET^18^ is a comprehensive platform integrating disease-associated genes information. It collects disease-associated genes from multiple sources: expert curated repositories, GWAS catalogues^17^, animal models and the scientific literature. As January 2019, it contains 628,685 gene-disease associations between 17,549 genes and 24,166 diseases/traits.

### Pharmacoepidemiologic validation

#### Study cohorts

We used the MarketScan Medicare Claims database from 2012 to 2017 for the pharmacoepidemiologic analysis. This dataset included individual-level procedure codes, diagnosis codes, and pharmacy claim data for 7.23 million patients. Pharmacy prescriptions of verapamil and amlodipine were identified by using RxNorm and National Drug Code (NDC).

#### Outcome measurement

For an individual exposed to the aforementioned drugs, a drug episode was defined as from drug initiation to drug discontinuation. Specifically, drug initiation was defined as the first day of drug supply (i.e. 1st prescription date). Drug discontinuation is defined as the last day of drug supply (i.e. last prescription date + days of supply) and without drug supply for the next 60 days. The fluticasone cohort included the first fluticasone episode for each individual, as well as the mometasone cohort. Further, we excluded observations that started within 180-day of insurance enrollment. For the extracted cohorts, demographic variables including age, gender and geographical location were collected. Additionally, diagnoses of hypertension (HT) and type 2 diabetes (T2D) (the International Classification of Disease [ICD] codes before drug initiation were collected, to address potential confounding biases. Last, a control cohort was selected from patients who were not exposed to fluticasone. Specifically, non-exposures were matched to the exposures (ratio 4:1) by initiation time of fluticasone, enrollment history, age and gender. The outcome defined by using ICD codes was time from drug initiation to diagnose of AD. For the fluticasone and mometasone cohorts, observations without diagnosis of AD were censored at the end of drug episodes. For the control cohort, the corresponding fluticasone episode starting date was used as the starting time. Observations without diagnosis of AD were censored at the corresponding fluticasone episode’s end date.

#### Propensity score estimation

We define NE = north east, NC = north central, S = south, W = west, T2D = type 2 diabetes, HT = hypertension and CAD = coronary artery disease. The propensity score of taking fluticasone vs. a comparator drug was estimated by the following logistic regression model:

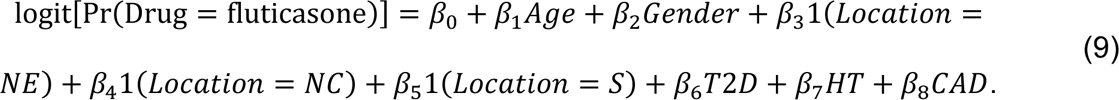

Further, among the subgroups defined by gender, the propensity score of taking verapamil vs. a comparator drug was estimated by the following logistic regression model:

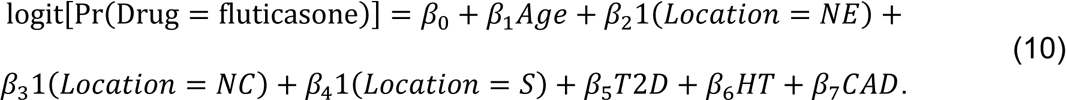

#### Statistical analysis

The survival curves for time to AD were estimated using a Kaplan-Meier estimator approach. We used the large number of covariates generated throughout the process to address clinical scenarios evaluated in each study. Additionally, propensity score stratified survival analyses were conducted to investigate the risk of AD between fluticasone users and non-fluticasone users, as well as fluticasone users and mometasone users. Specifically, for each comparison, the propensity score of taking fluticasone was estimated by using a logistic regression model, in which the covariates included age, gender, geographical location, T2D diagnosis and HT diagnosis. Further, propensity score stratified Cox-proportional hazards models were used to conduct statistical inference for the hazard ratios (HR) of developing AD between cohorts.

## Authors’ contributions

F.C. conceived the study. J.X., P.Z., Y.H., performed all experiments and data analysis.

L.B., J.L., C.C., L.L., A.A.P., J.B.L., and J.C. discussed and interpreted all results. J.X., F.C., Y.H. and J.C. wrote and all authors critically revised the manuscript and gave final approval.

---

**Supplementary Figure 1.**
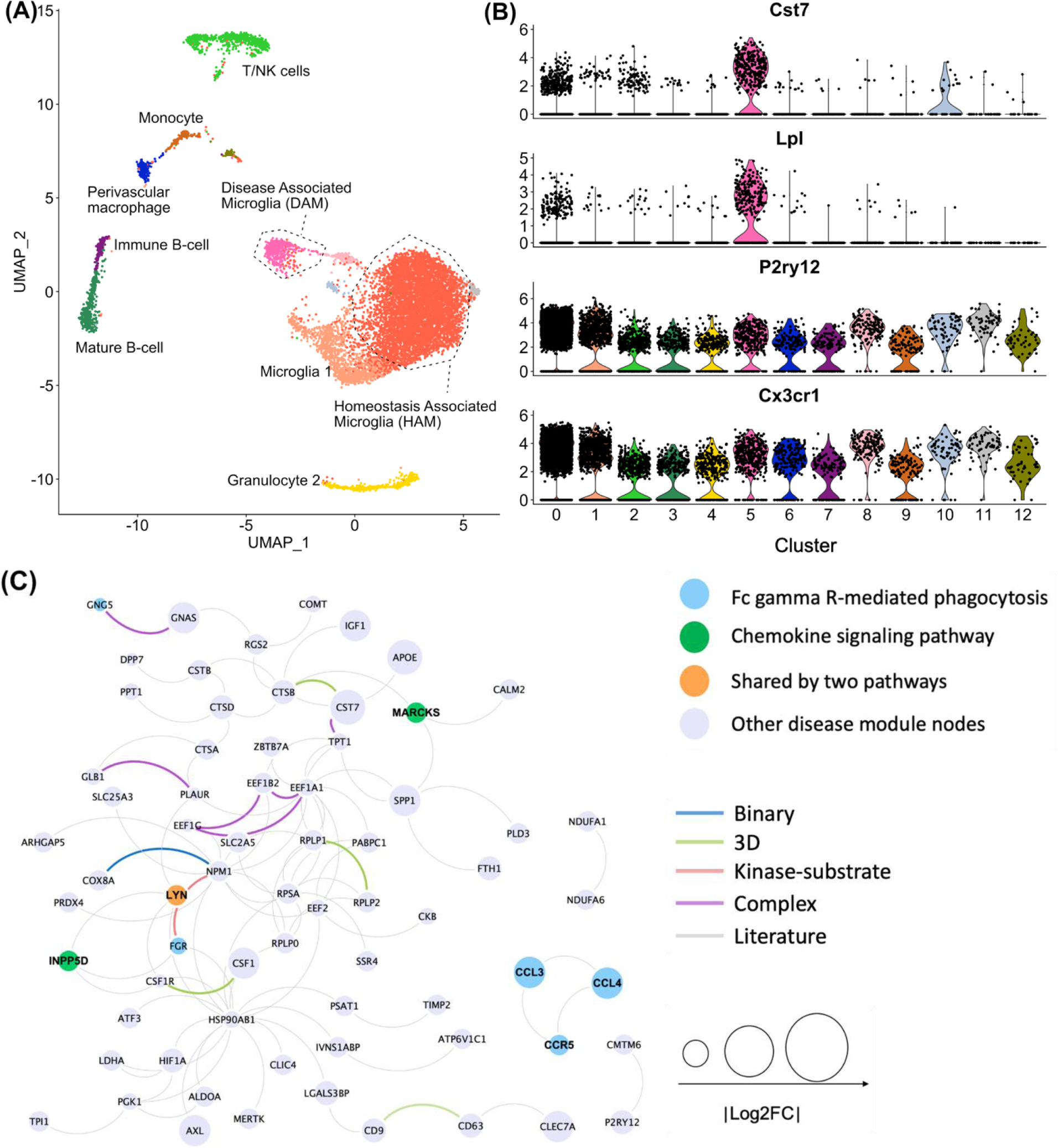
Discovery of disease-associated microglia (DAM) specific molecular networks using a scRNA-seq dataset (GSE98969). (**A**) UMAP plot of clustering 10,836 CD45+ cells into 13 sub-groups: gold cluster denoting the homeostasis associated microglia (HAM) and red cluster denoting the DAM. (**B**) Expression levels (stacked violin plots) of representative marker genes (up-regulation in DAM: *Cst7* and *Lpl* and down-regulation in DAM: *P2ry12* and *Cx3cr1*) in different microglia sub-clusters; (**C**) Extracted cell subtype DAM specific molecular network includes 69 nodes (proteins) and 97 edges (protein-protein interactions [PPIs]). Node sizes are proportional to their corresponding |log2FC|. Nodes are color coded by well-known KEGG immune pathways and edges are color coded by different experimental evidences of PPIs (**Methods**).

**Supplementary Figure 2.**
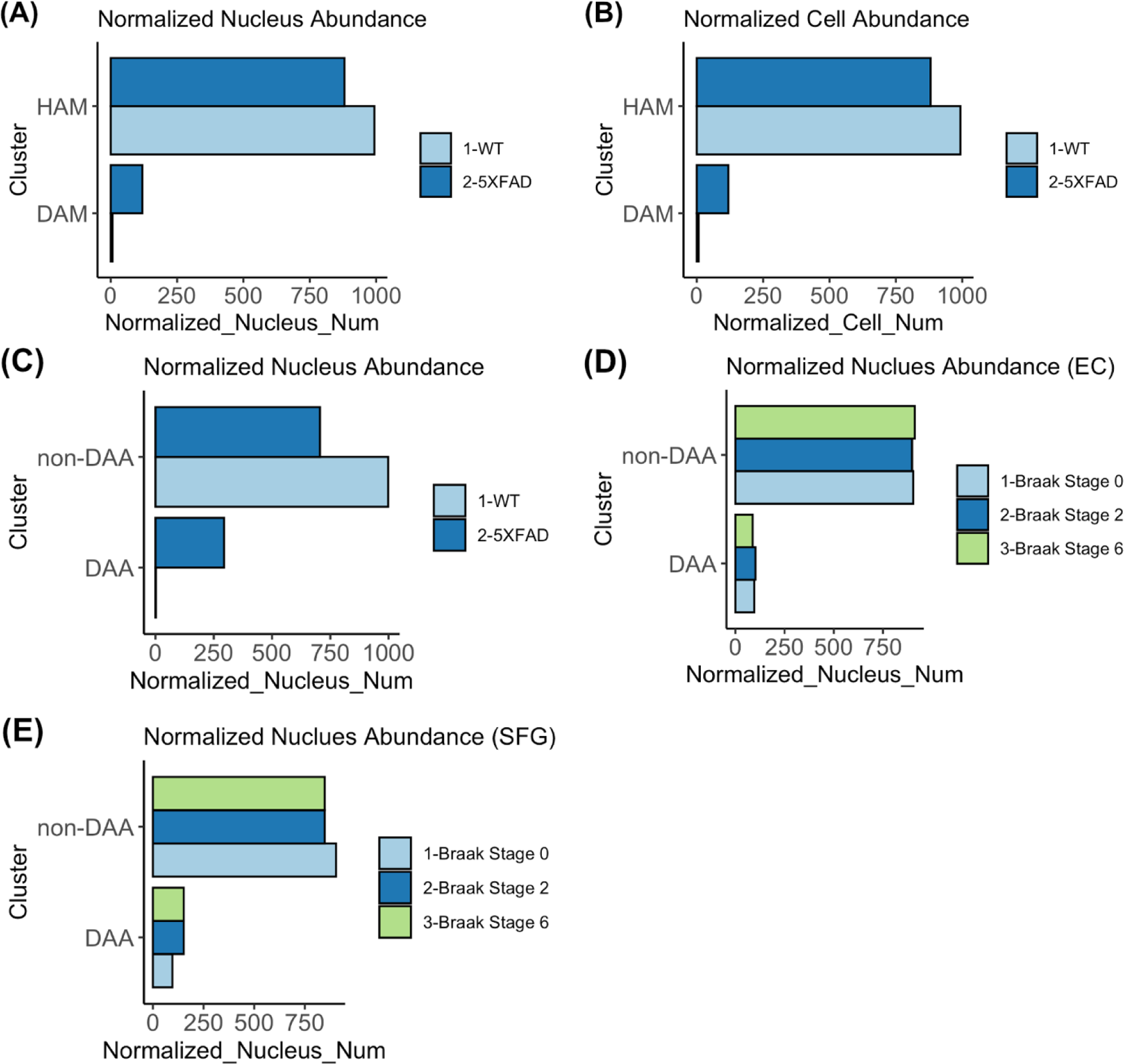
Nucleus / cell distribution in different immune cell subtypes. Normalized nucleus / cell abundance for homeostasis associated microglia (HAM) and disease associated microglia (DAM) clusters in both wild-type (WT) and 5XFAD mouse models (**A**) from sn-RNA seq dataset – GSE140511 and (**B**) from sc-RNA seq dataset – GSE98969. (**C**) Bar plot of normalized nucleus abundance in both disease associated astrocytes (DAAs) and non-DAA clusters considering both WT and 5XFAD mice. (**D** and Bar plot of normalized nucleus abundance in both DAA and non-DAA clusters considering human AD brains with Braak stages 0, 2, and 6 for brain region - (**D**) entorhinal cortex (EC) and (**E**) superior frontal gyrus. Detailed results are presented in Supplementary **Table 1**.

**Supplementary Figure 3.**
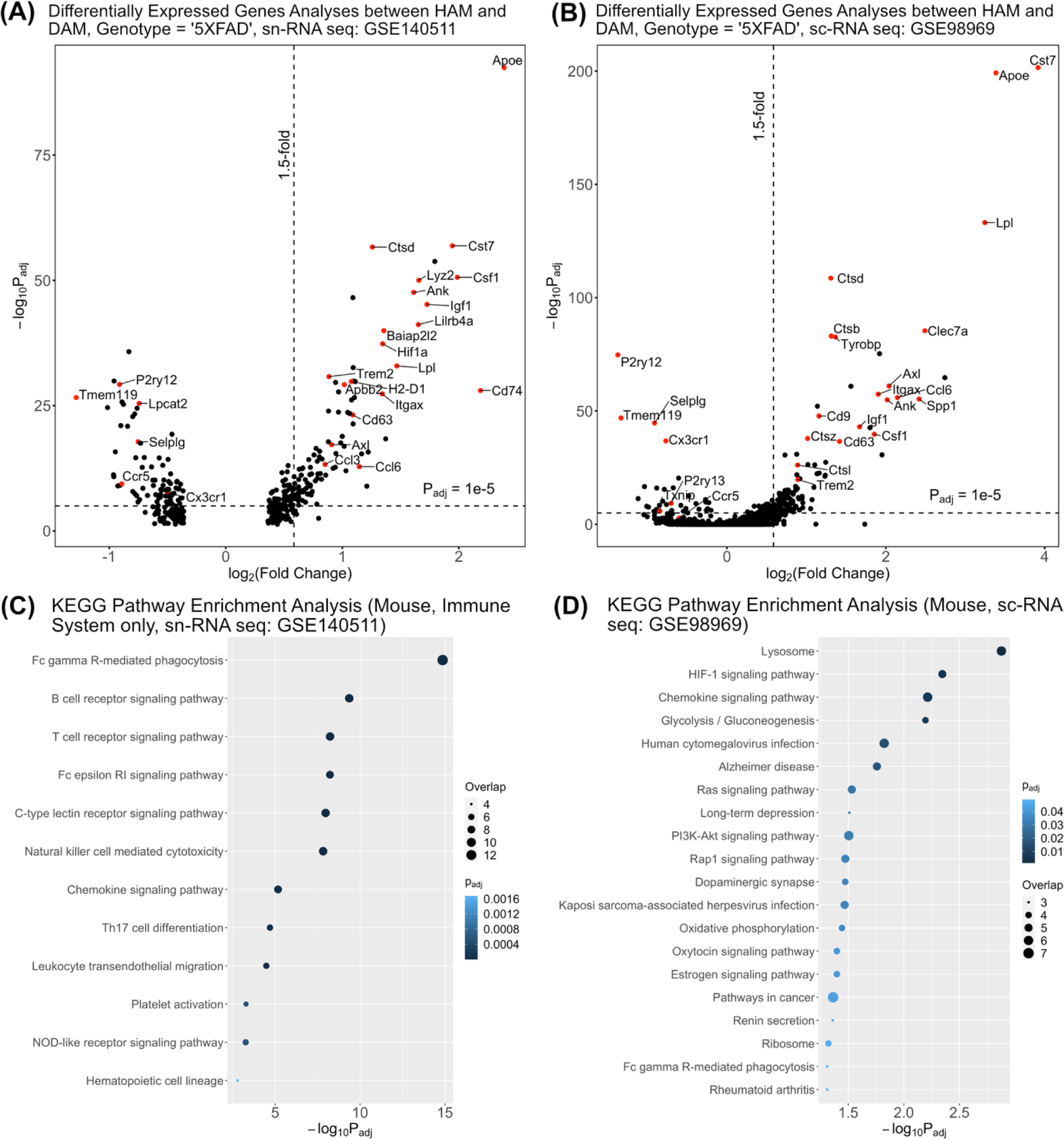
Differentially expressed genes and pathway enrichment analysis for disease associated microglia (DAM). Differential expressed gene analysis (volcano plot) between DAM and homeostasis associated microglia (HAM) with restricting to 5XFAD mice only using two different datasets: (**A**) GSE140511 (**Supplementary Table 1** and (**B**): GSE98969 (**Supplementary Table 2**). (**C**) Pathway enrichment analysis (**Supplementary Table 1**) for GSE140511. (**D**) Pathway enrichment analysis (**Supplementary Table 2**) for GSE98969.

**Supplementary Figure 4.**
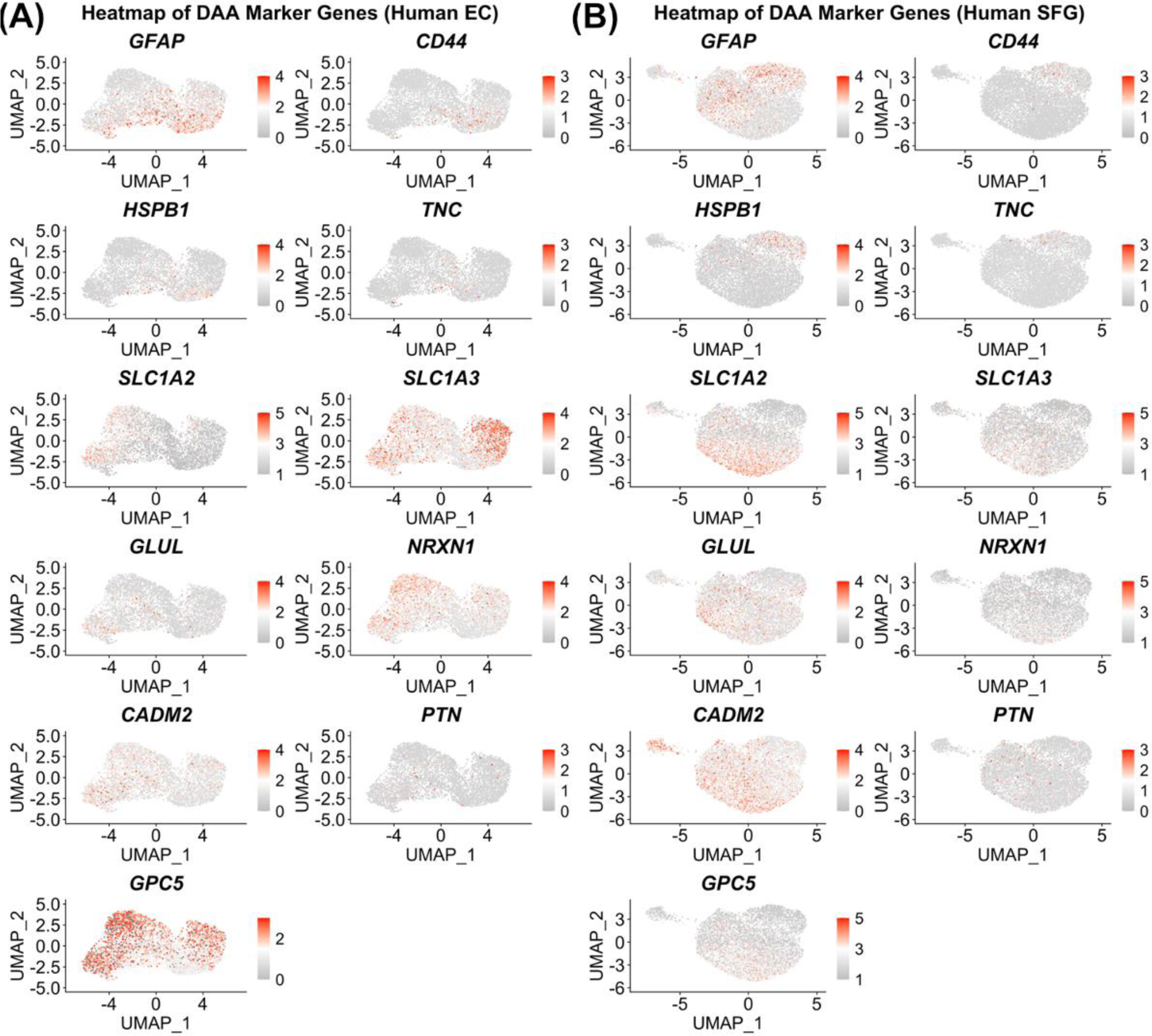
Expression levels (heatmap) of representative marker genes (up-regulation in DAA: *GFAP, CD44, HSPB1* and *TNC*, and down-regulation in DAA: *SLC1A2, SLC1A3, GLUL, NRXN1, CADM2, PTN* and *GPC5*) in all astrocyte sub-clusters, patients’ brain region: (**A**) entorhinal cortex (EC) and (**B**) superior frontal gyrus (SFG). Data source: GSE147528.

**Supplementary Figure 5.**
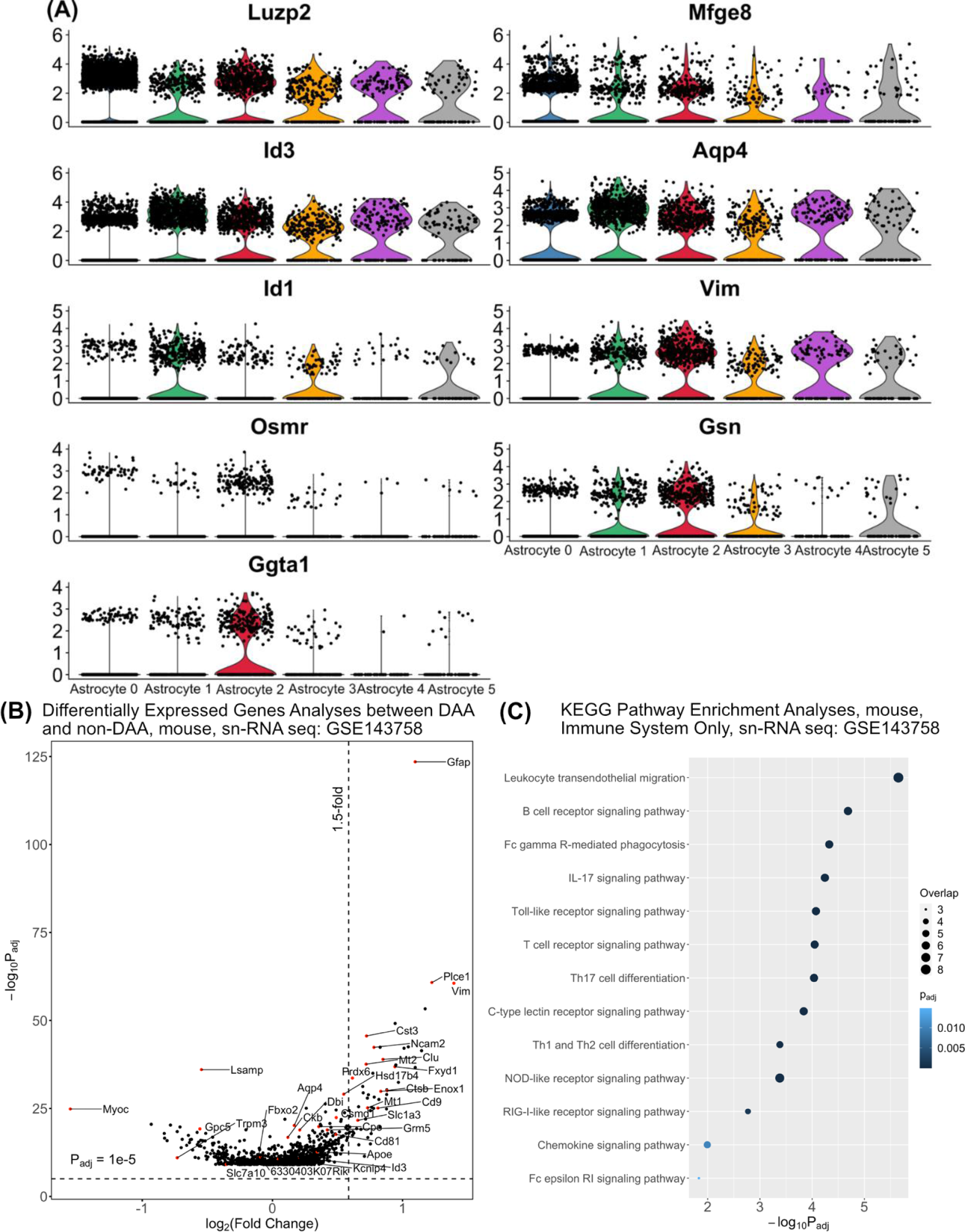
Differentially expressed genes and pathway enrichment analysis for disease associated astrocytes (DAAs) built from the AD transgenic mouse model (GSE143758). (**A**) Stacked violin plot displaying the expression patterns of 9 representative genes across different astrocyte sub-clusters. (**B**) Differential expressed gene analysis (volcano plot) between DAAs and non-disease associated astrocytes (non-DAAs) using 5XFAD mice. (**C**) Pathway enrichment analysis presented by 13 enriched KEGG immune system pathways (**Supplementary Table 3**).

**Supplementary Figure 6.**
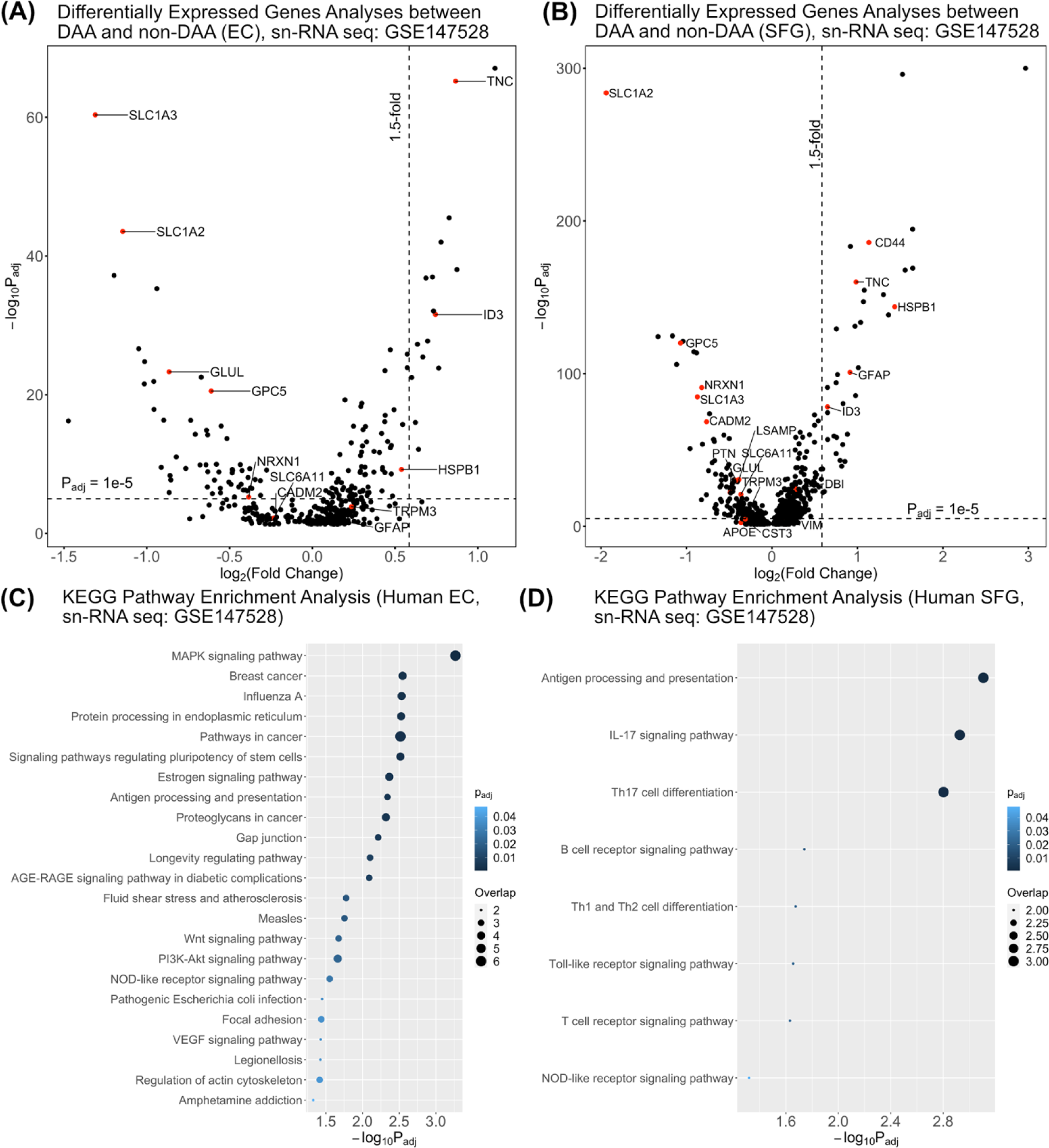
Differentially expressed genes and pathway enrichment analysis for disease associated astrocytes (DAAs) built from human AD patient snRNA-seq data (GSE147528). Differential expressed gene analysis (volcano plot) between DAAs and non-disease associated astrocytes (non-DAAs), patients’ brain regions: (**A**) entorhinal cortex (**B**) superior frontal gyrus; (**C**) Pathway enrichment analysis (see **Supplementary Table 4**) for molecular networks built from (**C**) entorhinal cortex (EC) and (**D**) superior frontal gyrus (SFG).

**Supplementary Figure 7.**
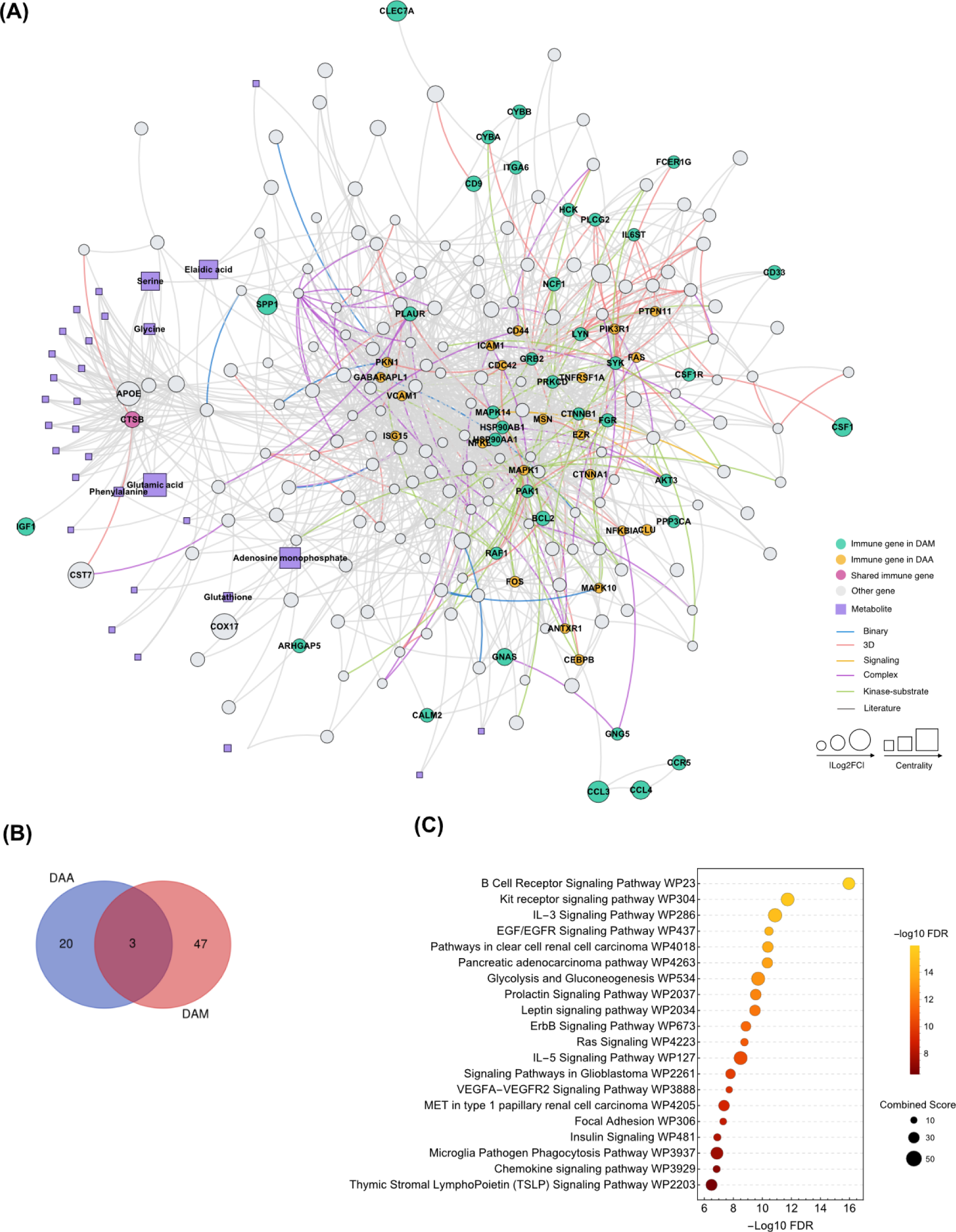
Network visualization and pathway enrichment analysis for disease associated astrocyte (DAA) and disease associated microglia (DAM). (**A**) A module illustrating the network-based relationship between DAA and DAA immune genes associated with AD-related metabolites. (**B**) Venn diagram of enzymes from DAA and DAM. (C) Pathway enrichment of 70 enzymes in DAA and DAM.

**Supplementary Figure 8.**
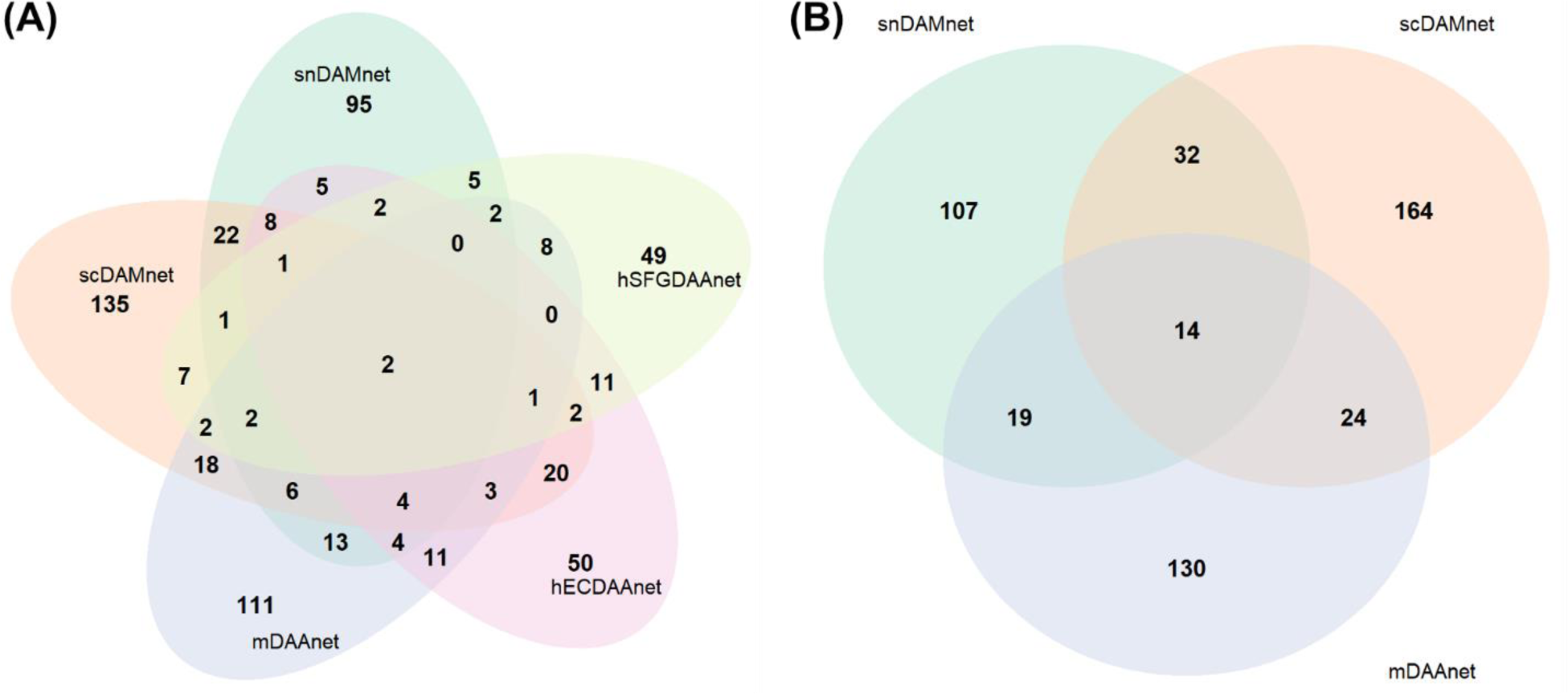
Network-based discovery of drug candidates. Venn diagrams show the relations of potential drug candidates predicted among (**A**) all datasets and (**B**) 3 mouse model datasets. To be specific, snDAMnet is a molecular network based on snRNA-seq mouse model dataset – GSE140511, scDAMnet is a molecular network built from scRNA-seq mouse model dataset – GSE98969, mDAAnet is a molecular network built from snRNA-seq mouse model dataset – GSE143758 and hECDAAnet and hSFGDAAnet are molecular networks based on snRNA-seq human AD brain dataset – GSE147528.

